# Superabsorbent crosslinked bacterial cellulose biomaterials for chronic wound dressings

**DOI:** 10.1101/2020.03.04.975003

**Authors:** Daria Ciecholewska-Juśko, Anna Żywicka, Adam Junka, Radosław Drozd, Peter Sobolewski, Paweł Migdał, Urszula Kowalska, Monika Toporkiewicz, Karol Fijałkowski

**Author notes:** **Corresponding author** Karol Fijałkowski, Department of Microbiology and Biotechnology, Faculty of Biotechnology and Animal Husbandry, West Pomeranian University of Technology, Szczecin, Piastów 45, 70-311 Szczecin, Poland.

## Abstract

In this work, we present novel *ex situ* modification of bacterial cellulose (BC) polymer, that significantly improves its ability to absorb water after drying. The method involves a single inexpensive and easy-to-perform process of BC crosslinking, using citric acid along with catalysts, such as disodium phosphate, sodium bicarbonate, ammonium bicarbonate or their mixtures. In particular, the mixture of disodium phosphate and sodium bicarbonate was the most promising, yielding significantly greater water capacity (over 5 times higher as compared to the unmodified BC) and slower water release (over 6 times as compared to the unmodified BC). Further, our optimized crosslinked BC had over 1.5x higher water capacity than modern commercial dressings dedicated to highly exuding wounds, while exhibiting no cytotoxic effects against fibroblast cell line L929 *in vitro*. Therefore, our novel BC biomaterial may find application in super-absorbent dressings, designed for chronic wounds with imbalanced moisture level.

## 1. Introduction

A wide spectrum of health issues, frequently referred to as “lifestyle diseases,” poses an increasing threat to modern western societies, due to their serious health complications (1). In particular, chronic wounds are among the most persistent and serious, because they do not follow the natural trajectory of healing and, if infected, threaten the life of the patient (2). Due to their persistence, chronic wound treatment is a significant burden for patients, their families, and health-care professionals, as well as healthcare systems in general (3). Therefore, prevention and treatment of chronic wounds is one of the most important challenges of contemporary medicine. To date, a number of approaches to managing chronic wounds have been developed, including T.I.M.E. (Tissue, Infection, Moisture, Epithelialization), B.B.W.C. (Biofilm-Based Wound Care), and W.A.R. (Wounds at Risk) (4,5,6). Although these strategies address the process of wound healing from different angles, they share a common denominator: maintenance of proper moisture level. In western medicine, keeping wounds dry was the conventional wisdom until landmark work by George Winter in 1962, who unquestionably demonstrated that moist wounds heal better than dry ones (7). Winter’s paper became the basis for the moist wound-healing concept, which is generally accepted and has been further developed ever since.

The intrinsic moisture of a wound is provided by a fluid referred to as “exudate” (8). In acute wounds, exudate accelerates the healing process, while in chronic wounds both over- or under-production of exudate can have a negative effect. Therefore, maintenance of the proper amount of exudate in chronic wounds is crucial for healing (9).

Additionally, to provide the appropriate healing conditions, the wound itself should be protected from microbial contamination and other external factors. Presently, there is a wide range of modern dressings dedicated to protecting either dry, low-exuding or highly exuding wounds (10–13). Nonetheless, there is still an ongoing search for a dressing material that would yield quick closure of chronic wounds regardless of type and amount of exudate.

One of the candidates for such an ideal dressing material is bacterial cellulose (BC). This polymer is biosynthesized primarily by acetic acid bacteria (14), of which Gram-negative *Komagataeibacter xylinus* is the most effective producer (15). The high purity, crystallinity and density, binding capacity, shape, and water retention (16–18) of BC has drawn extensive interest of research teams from all over the world. This is reflected by the rapidly increasing number of scientific reports on BC (19), particularly over the past 15 years, investigating its wide range of potential applications in many branches of industry (20). In the biomedical field, this multifaceted material has already been used, among others, in drug-delivery carriers (21) and in wound dressings for patients with extensive burns (16). These applications are possible thanks to the structure of BC, consisting of thin, loosely arranged fibrils, separated by “empty” spaces (22). This affects the liquid capacity of BC, which depends primarily on fibril arrangement, surface area, and porosity. Overall, the more space is available between the BC fibrils, the more liquid penetrates and adsorbs into the material. It has been estimated that approx. 97-98% of total BC weight consists of water (23,24).

However, from the point of view of industry, keeping materials dry during storage is important for several obvious reasons, including prolonging the product’s expiration date and microbiological safety. While BC can be subjected to dehydration, the evaporation of water from the interfibrillar spaces causes the collapse of the BC structure (22,25). This irreversible process deprives BC of its most unique properties. After dehydration, the three-dimensional polymer network is unable to regenerate and its ability to re-swell and absorb liquid is significantly reduced, as compared to never-dried cellulose (26,27). Therefore, dried BC is less useful within the scope of biomedical applications (22,28). For this reason, several *ex situ* modifications have been proposed to counteract the disadvantageous consequences of BC dehydration, for example the production of BC-fibrin or BC-gelatin composites, decreasing BC crystallinity, or freeze drying (28–32). Crosslinking is another type of *ex situ* modification method, typically obtained by linking at least two hydroxyl groups of single cellulose molecule or two or more hydroxyl groups of adjacent cellulose molecules. As a result, the material becomes stiffer and preserves its three-dimensional structure (33–35). Crosslinking efficiency can be further increased by using compounds that act as catalysts (CATs) for the reaction (36).

Crosslinking reactions have been used to modify many biopolymers; however, to date only a single scientific publication by Meftahi et al. (37) reports the use of crosslinking to modify cellulose produced by microorganisms. In their work, Meftahi et al. (37) reported that the use of citric acid (CA) as a crosslinking reagent, with sodium hypophosphite (SHP) catalyst (CAT), resulted in significantly enhanced (up to 3 times) levels of re-hydration of previously dried BC. CA and SHP have also already been used in a number of polymer studies involving crosslinking of hydrogels, plant cellulosic materials, and bio-composites (37–41).

CA is a particularly promising crosslinker not only because of its safety and non-toxicity, but also because of the stable crosslinking bonds it forms with cellulose. Importantly, CA is easily accessible and inexpensive, making this reagent commercially attractive. On the other hand, the use of SHP in chemical reactions can be problematic, because thermal SHP decomposition results in the formation of phosphine, a highly toxic and extremely flammable gas (42). Further, SHP itself can have a negative environmental impact, because leached hydrogen phosphide compounds can contaminate surface waters (43).

In the present work, our primary goal was to obtain dry BC, displaying very high water-related parameters, which can be used as a super-absorbent dressing for chronic wounds. Towards this aim, we crosslinked BC with CA and tested a range of compounds as catalysts, the majority of which have never been used for this purpose. Importantly, focusing on biomedical applications and safety, we selected reagents that are all on the list of substances generally recognized as safe (GRAS), according to the U.S. Food and Drug Administration and are widely used in the food industry, for example as chemical leavening in baking (44,45). This fact served as the basis for our study: these compounds release large amount of gases during reactions with acids and as a result of thermal decomposition. We hypothesized that released gas would fill the spaces between the crosslinked fibrils and BC layers, preventing them from collapsing during dehydration. Further, we assumed that the released gases could form new cavities within the BC matrix, additionally increasing water sorption capacity. Finally, in order to facilitate future translation and commercialization, we also assessed the specific CATs from the point of view of their potential impact on the environment, cytotoxicity, and cost-efficiency.

## 2. Materials and methods

### 2.1. Bacterial cellulose synthesis and purification

To obtain BC in a form of pellicles of consistent shape and diameter, the cellulose-producing *Komagataeibacter xylinus* strain (American Type Culture Collection – ATCC 53524) was cultured in 50 mL plastic tubes (Becton Dickinson and Company, USA), 24- or 96-well cell culture plates (Nest Scientific USA Inc., USA) under stationary conditions using Hestrin-Schramm (H-S) culture medium, over 7 days at 28°C. Obtained BC samples were purified from bacteria and culture medium components using 0.1 M sodium hydroxide solution at 80°C for 90 min, followed by rinsing with distilled water until the pH was neutral. The diameter of the BC pellicles obtained in plastic tubes was 2.5 cm, the average thickness was 0.9 cm, and the average weight was 4.5 g, whereas the diameter, average thickness and weight of BC pellicles obtained from 24- and 96-well plates were 1.25 cm, 0.4 cm, and 0.9 g and 0.6 cm, 0.15 cm and 0.3 g, respectively.

### 2.2. Crosslinking procedure

The crosslinking reaction was carried out using aqueous solutions of CA and different CATs. Six substances were used as CATs: disodium phosphate (later referred to as CAT1), sodium bicarbonate (later referred to as CAT2), disodium phosphate and sodium bicarbonate (1:1 mass ratio, later referred to as CAT3), ammonium bicarbonate (later referred to as CAT4), disodium phosphate and ammonium bicarbonate (1:1 mass ratio, later referred to as CAT5), sodium bicarbonate and ammonium bicarbonate (1:1 mass ratio, later referred to as the CAT6). Additionally, sodium hypophosphite (later referred to as CAT7) was used as a reference, because it is well described in the literature related to crosslinking of plant and also bacterial cellulose (37, 46–48). All of the chemicals were purchased from POCH, Poland.

The crosslinking procedure consisted of four stages: ***i)*** impregnation of BC with CA and CAT for 24 h at 28°C, ***ii)*** incubation at high temperature to induce the crosslinking reaction, ***iii)*** purification with distilled water to remove unbound CA molecules and CATs, ***iv)*** drying at room temperature (**Figure 1**).

**Figure 1.**
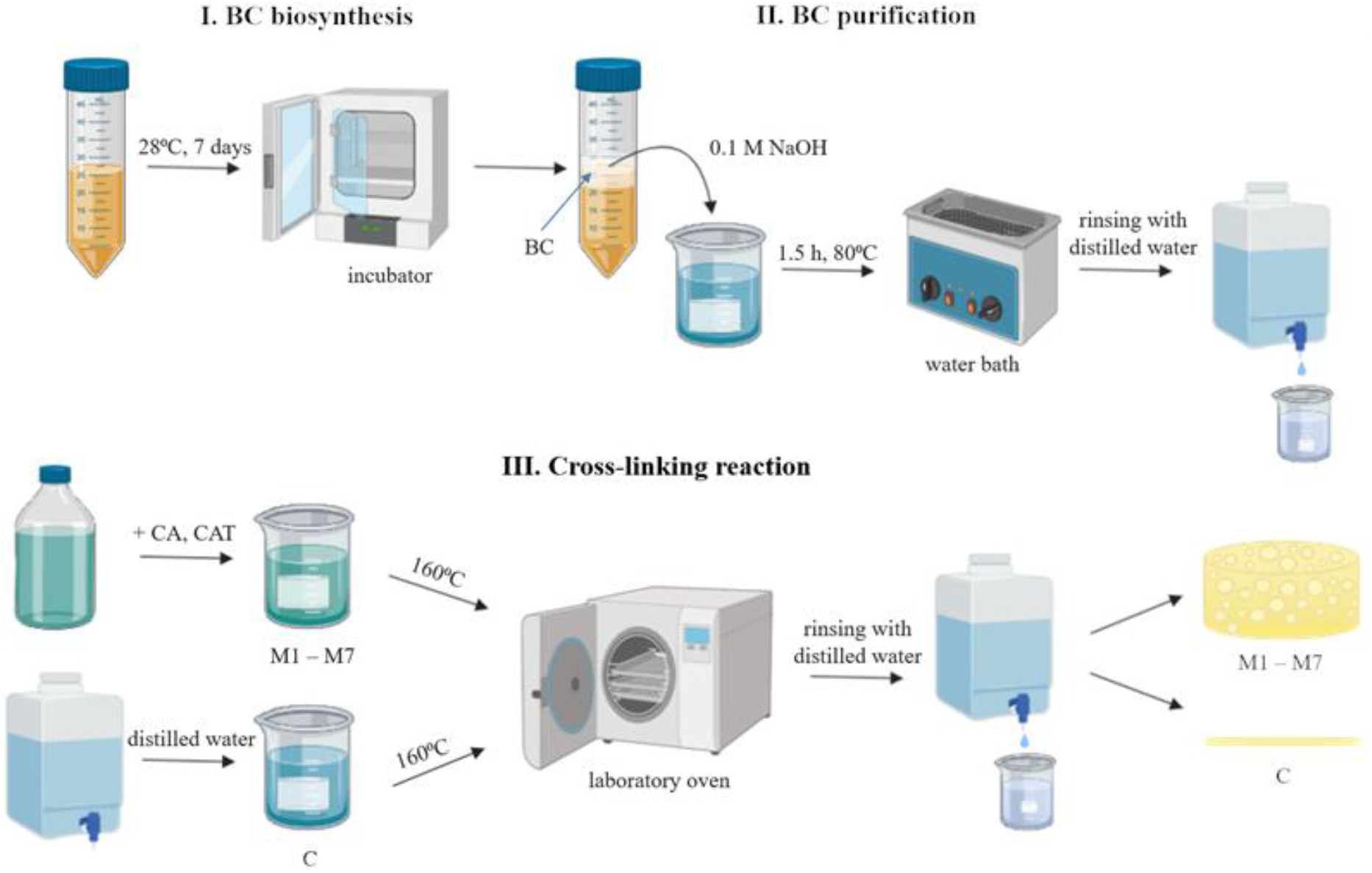
BC modification procedure (created by BioRender.com). C – control BC sample; CA – citric acid; CAT – catalyst; M1-M7 – modified BC samples.

The BC samples obtained as a result of aforementioned modifications were referred as M1-M7, depending on the CAT used (CAT1-CAT7, respectively). As a control, BC treated with just distilled water, without CA or CAT, was used (later referred to as C).

#### Optimization of the BC crosslinking reactions

The first parameters to be optimized were the concentrations of CA and CATs. The following concentrations (in distilled water, m/v) were tested: CA – 30%, 20%, 10%, 5%; CAT – 15%, 10%, 5%, 2.5%. Additionally, the reaction using 20% CA without any CAT was performed as a comparative control setting. The initial CA to CAT mass ratio (2:1) and reaction temperature (160°C) were selected based on the previously published conditions (47–49). The reaction was carried out until BC was completely dehydrated, that is until the weight of the sample stabilized.

Following the determination of the optimal concentrations of CA and CATs, we optimized the remaining parameters: CA to CAT mass ratio, reaction time and temperature. The following mass ratios between defined optimal concentrations of CA and CAT were tested: 1:1, 1.5:1, 2:1, 2.5:1, 3:1, 1:2. The role of crosslinking reaction temperature was assessed in the range between 120 and 180 °C and the following reaction times: 5, 15, 30, 60, 75 and 90 min. Further analyzes, presented below, were carried out for BC crosslinked in the optimal conditions.

### 2.3. Evaluation of the properties of BC crosslinked under optimal conditions using various catalysts

#### 2.3.1. Analysis of the BC macrostructure – stereoscopic microscope

The macromorphological structure of modified and unmodified (control) BC samples was evaluated using stereoscopic microscope (Leica S9i, Leica Microsystems, Wetzlar, Germany). The analyzes were performed for dry and wet (rehydrated) BC samples. Both, the surface and the layer structure visible in cross-section were assessed.

#### 2.3.2. Analysis of the BC microstructure – scanning electron microscope

Modified BC samples were fixed in glutaraldehyde (POCH, Poland) for 0.5 h and washed in the following concentrations of ethanol (POCH, Poland): 10%, 25%, 50%, 60%, 70%, 80%, 90%, 100%, respectively, 5 min at each concentration. Subsequently, the BC was subjected to sputtering with Au/Pd (60:40) using EM ACE600, Leica sputter (Leica Microsystems, Wetzlar, Germany). Both the surface and the layer structure visible in cross-section were assessed. The surfaces of the sputtered samples were examined using Auriga 60 scanning electron microscope (SEM, Auriga 60, Zeiss, Oberkochen, Germany), whereas cross sections of the samples were analyzed using VEGA3 scanning electron microscope (VEGA3, Tescan, Brno, Czech Republic). The BC microfibril diameters were analyzed using software integrated with Auriga 60 SEM, while micro-pore size was analyzed using ImageJ software (50).

#### 2.3.3. Elemental analysis of BC surface – energy-dispersive X-ray spectroscopy

Samples were placed on aluminum tables, sputtered with gold (20 nm) using EM ACE600, Leica sputter (Leica Microsystems, Wetzlar, Germany), and placed in the scanning electron microscope chamber (Auriga 60, Zeiss, Oberkochen, Germany). An X-ray microanalysis (EDX) was carried out using an EDX Oxford detector (Oxford Instruments, Abingdon, United Kingdom). The working distance was 5 mm and the voltage 20 kV. For characterization of sample composition, the AZtec system (Oxford Instruments, Abingdon, United Kingdom) was used.

#### 2.3.4. Determination of chemical composition of the BC – attenuated total reflectance Fourier transform infrared (ATR-FTIR) spectral studies

The assessment of the chemical composition of the BC samples was performed using a Bruker Alpha FTIR spectrometer with an ATR adapter. Spectra (32 scans) were collected over wavenumber range of 4000-400 cm^−1^ with a resolution of 4 cm^−1^. The obtained ATR-FTIR spectra were analyzed using the Spectragryph 1.2 software. Additionally, the 2DShige© v1.3 program was used to preform 2D correlation analysis. 2D correlation spectra were visualized using OriginPro 8 software (Origin Software Solutions, United Kingdom) (51,52).

The chemometric principal component analysis (PCA) of all spectra was performed with using R software and the *FactoMineR* and *Factorextra* packages via RStudio software (53). Prior to the analysis, the spectra in the of range 1850 cm^−1^ to 850 cm^−1^ were normalized to the peak area at 1200 cm^−1^. The areas of integrated peaks were used as input data for PCA.

#### 2.3.5. Determination of water swelling ratio

To determine the swelling ratio (SR (%)), BC pellicles were first dried at room temperature to completely remove water content. The dry BC samples were then immersed in distilled water and weighed – first after 1 min, and then every 10 min up to 60 min. After 60 min, the BC samples were left in water and their weight was measured after 24 h. The percentage of water absorption was calculated using the Equation (1) (54):

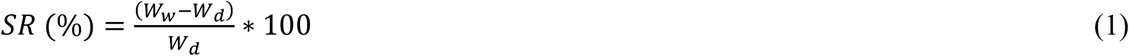

where, W_w_ is the weight of the swollen BC and Wd is the dry weight of the sample.

#### 2.3.6. Determination of water holding capacity

Dry BC pellicles were weighted, immersed in distilled water for 24 h to obtain maximum absorption level, and then weighed again. The ability to hold water was determined using two methods. First, BC samples were arranged on cell strainers placed in 50 mL tubes, centrifuged (5804R, Eppendorf, Germany) at 200 *g*, and then weighted every 10 min for 100 min. In the other method, BC samples were placed in an incubator at 37°C and weighted every 10 min for 240 min. Water holding capacity (WHC (%)) was calculated using the Equation (2):

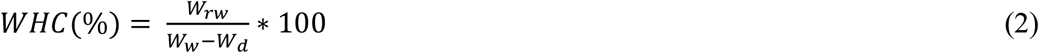

where, W_rw_ is the weight of water retained in BC during drying/centrifuging, W_w_ is the initial weight of wet BC, and Wd is the dry weight of the sample.

#### 2.3.7. Determination of the BC density

The density of the BC samples was determined using hydrostatic balance (XA 52/Y, Radwag, Poland) in methanol as a standard liquid. The weight of samples was measured at room temperature in air, as well as in methanol. The sample density was determined using the Equation (3) (55):

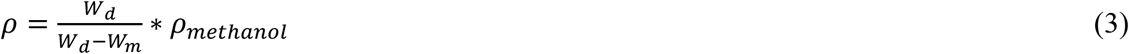

where, *W*_*d*_ is the weight of the dry sample in the air, *W*_*m*_ is the weight of the sample in methanol, and ρ_methanol_ is the density of methanol.

### 2.4. Evaluation of the crosslinking efficiency under optimal conditions using various catalysts

The BC crosslinking efficiency under optimal, defined reaction conditions was assessed as the BC weight percent gain (WPG (%)) compared to the weight of unmodified BC (control) and calculated according to the Equation (4) (56):

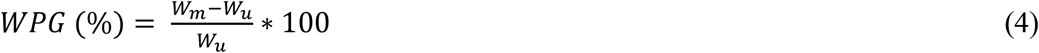

where: W_m_ is the dry weight of modified BC, and W_u_ is the dry weight of unmodified BC.

### 2.5. Determination of the cytotoxicity of BC crosslinked under optimal conditions using various catalysts

#### 2.5.1. Direct contact assay

All cell culture reagents and cell lines were purchased from Sigma-Aldrich (Poznań, Poland). For direct contact assay based on the ISO 10993-5:2009 standard (59), 1 × 10^4^ L929 cells (mouse fibroblasts) were seeded per well in a 96-well plate. Following 24 h of culture in Dulbecco’s Modified Eagle Medium (DMEM) containing 10% fetal bovine serum (FBS), 2 mM L glutamine, 100 U/mL penicillin, and 100 μg/mL streptomycin (hereafter referred to as “complete growth media”) in an incubator (5% CO_2_, 37°C), the cells were assessed to be ~50% confluent, the medium was aspirated, and previously autoclaved modified BC pellicles with a diameter of 6 mm were placed on top of the cell layer. Each disc was pretreated by soaking in 2 mL of complete growth medium for 1 h. CellCrown tissue culture inserts were then placed on top of the discs to prevent them from floating and media was replaced. As a sham control, CellCrown tissue culture insert was placed directly into the well. After another 24 h of culture in 5% CO_2_ at 37°C, wells were examined using inverted light microscope (Delta Optical IB-100, Poland) and then resazurin stock (0.15 mg/mL) was added to each well (1:6 dilution) (57). After 4 h of incubation the viability was assessed using fluorescent plate reader (Synergy HTX, Biotek, USA) at wavelengths of 540 nm excitation and 590 nm emission. Complete growth media in an empty well was used as a blank. The results were expressed as percent of cell viability, and calculated by the Equation (5) (58):

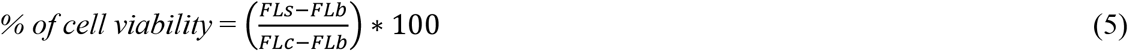

where FL is the fluorescence intensity (arbitrary units) and indexes *s, b,* and *c* refer to sample, blank, and control, respectively.

#### 2.5.2. Extract assay

Extracts of crosslinked BC were prepared according to the ISO 10993-5:2009 standard (59). Briefly, for each tested material 8 BC pellicles were placed in a 24-well plate well and covered with 1.5 mL of complete growth medium. As a sham control, 1.5 mL of media was pipetted into an empty well. The plate was then placed into 37°C CO_2_ incubator for 24 h of extraction. Simultaneously, 1 × 10^4^ L929 cells were seeded per well in a 96-well plate. Following 24 h of culture in 5% CO_2_ at 37°C, the cells were assessed to be ~50% confluent, the media was aspirated and replaced with 100 μL of extract – 5 technical replicates were performed per sample. After another 24 h of culture, cells were examined using inverted light microscope and then cell viability was assessed as described above.

#### 2.5.3. Confocal microscopy analysis of fibroblast viability on BC pellicles

L929 fibroblasts were cultured on BC samples for 5 days, followed by fixation in 3.7% formaldehyde solution and staining with SYTO-9 and Propidium Iodide (PI) dyes (a mix of both from ThermoFisher) to visualize DNA in live and dead cells, respectively. The samples were then imaged using an upright Leica SP8 confocal microscope (Leica Microsystems, Wetzlar, Germany). Stacks of confocal 8-bit images with a voxel size of 0.465 × 0.465 × 1.5 μm were acquired using a dry 20x objective (NA 0.75) with the pinhole was set to 1 AU. SYTO-9 was excited with 488 nm laser line and 492-526 nm emission range was collected, while PI fluorescence was excited with 552 nm laser line and 561-611 nm emission range was recorded. The presence of cellulose surface below cultured cells was confirmed in reflection mode using a 638 nm laser line. The acquisition was performed in sequential mode. Three-dimensional rendering was performed in Imaris software (Bitplane, United Kingdom).

### 2.6. Statistical analysis

Data are shown as means ± standard errors of the means (SEM) obtained from at least three different measurements (plus technical repetitions). Statistical differences between different BC samples were determined by one-way analysis of variance (ANOVA) and Tukey’s post hoc test. All analyses were considered statistically significant when the P value was less than 0.05. Statistical tests were conducted using Statistica 9.0 (StatSoft, Poland).

## 3. Results and Discussion

### 3.1. Optimization and determination of the BC crosslinking parameters

The primary aim of this work was to obtain super-absorbent biomaterial. Therefore, the water-swelling ratio (SR (%)) was used as the primary parameter to assess the effectiveness of each BC modification.

The SR (%) values for BC samples modified using different CATs under the same reaction conditions displayed a comparable trend. The most beneficial (for BC sorption properties) set of parameters for the crosslinking reaction was: 20% CA concentration with 10% of the appropriate CAT, 2:1 mass ratio of CA:CAT, reaction temperature of 160°C, and reaction time of 75 min. The SR (%) results for entire data set of the results from the optimization of crosslinking process are presented in the Supporting Information (**Figure S1**, **Tables S1-S9**).

### 3.2. Macro and micromorphological characteristics of modified BC

Macromorphological examination demonstrated that dried, modified BCs, contrary to the unmodified samples, possess an arranged and multilayer structure regardless of CAT used (**Figure 2**, **Figure S2**).

**Figure 2.**
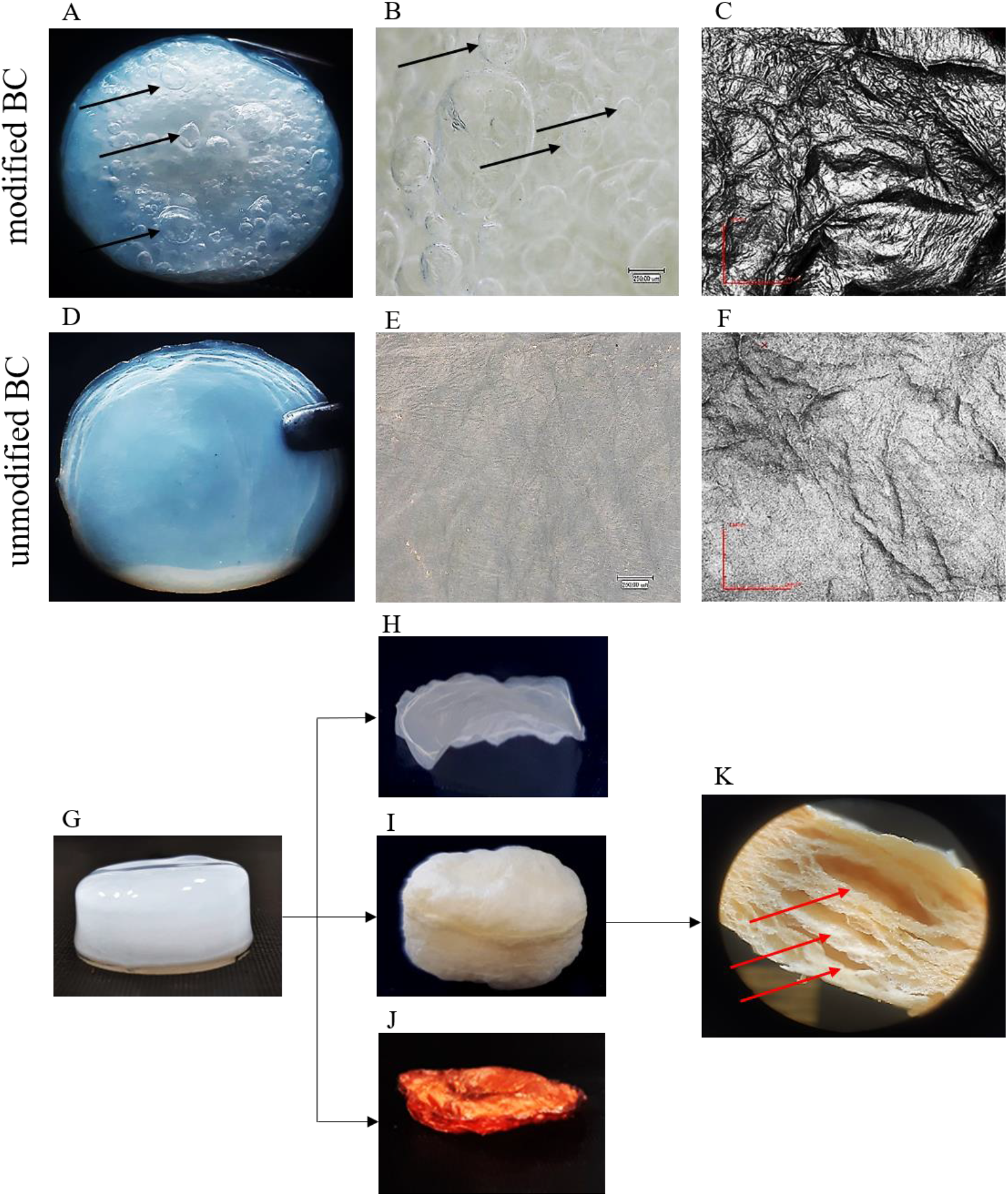
The macromorphological differences in modified and unmodified BC. A,B – top view of wet modified BC; C – surface of dry modified BC; D,E – top view of unmodified wet BC; F – surface of unmodified dry BC; G – side view of unmodified never-dried BC; H – side view of unmodified dry BC; I – side view of dry modified BC; J – side view of dry BC treated with 20% CA w/o CAT; K – cross-section of dry modified BC. Pictures of modified BC are presented on the example of M3 (BC modified with 20% CA + 10% CAT3). The pictures of cross section and surface of all modified BC samples are presented in Supporting Information (**Figures S2-S4**). Black arrows (A,B) indicate air bubbles on the surface of the sample, whereas red arrows (K) indicate spaces between BC layers.

This multilayer spatial arrangement of BC is directly related to the biological and physical processes of its polymer synthesis (14). Cellulose fibrils secreted by the bacterial cells into the extracellular space form a thin layer on the surface of the medium (60). Over a time, a new cellulose layer appears on the surface, pushing the older one down and this process repeats until depletion of oxygen and nutrients in medium. The fact we observed multilayers in our crosslinked BC samples confirmed that the process successfully prevents the collapse of the 3D structure during drying.

Cross sections of the dried modified BC samples revealed the presence of empty spaces between the layers in all of the modified BC pellicles, but not in the case of unmodified BCs and BCs modified only with CA (**Figure 2G-K**, **Figure S2**). This provides strong evidence in support of our hypothesis that gas bubble formation occurs within the BC structure during crosslinking reaction with use of the tested thermo-sensitive catalysts (**Figure 2A**, **Figure 2B**, **Figure S3**). Also, it is worth mentioning that the surface of the modified BC was also heavily folded, in contrast to the relatively smooth surface of unmodified BC, but no specific pattern of surface folding with different CATs was observed (**Figure 2A-F**, **Figure S3**, **Figure S4**). Thus, the results of macromorphological examination help to explain why modified BC samples, particularly these crosslinked with addition of CAT3, displayed significantly higher SR (%) than unmodified BC (see **Figure S1**, **Tables S1-S9**).

In order to assess the microscale architecture of crosslinked BC, we used SEM imaging. Overall, SEM images of the surfaces of BC pellicles demonstrated that the micro-porosity and BC layer structure were preserved in all of the studied reaction conditions (**Figure S5**). Fibril thickness and average pore diameter were comparable between all modified BC pellicles (regardless of CAT used) and control BC samples (**Figures S6-S8**). In contrast, the SEM imaging of BC cross sections showed that morphology of stratification, as well as the formation of spaces between layers was CAT-specific (**Figure 3**). In all modified BC pellicles, multiple spaces appeared as a result of delamination or blistering. However, we did not observe such stratification patterns in the case of control BC, which was characterized by a typical homogeneous structure throughout the entire section.

**Figure 3.**
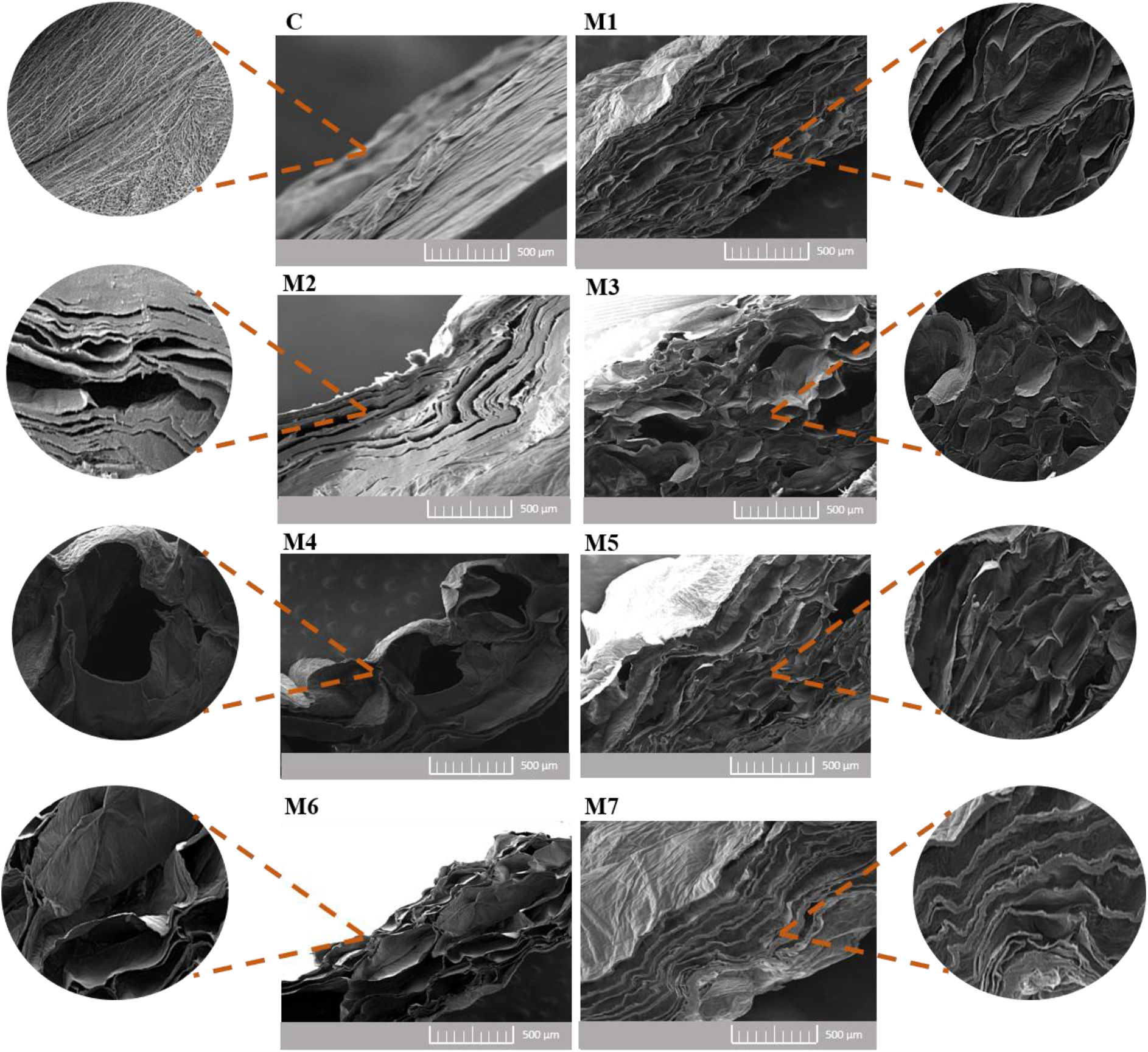
Cross sections of BC pellicles visualized by scanning electron microscopy (SEM, VEGA3, Tescan, Brno, Czech Republic) at 500x and 2000x magnification for images in the center and on the edges of the figure, respectively. C – control BC sample; M1-M7 – modified BC samples.

For the case of M1, SEM imaging of cross sections demonstrated regular corrugations of the matrix, the presence of numerous, but relatively small, regular bubbles (air spaces), and moderate layer stratification (**Figure 3**). In contrast, M2 was characterized by greater degree of corrugation. Instead of the bubbles present in M1, longitudinal air spaces were observed between the separate layers of M2. The structure of M3 resembled M1, except that the degree of folding was greater and the bubbles (visible as black spots) were larger and more deeply located. Meanwhile, M4 microstructure was characterized by a very high degree of folding and the presence of very large air spaces taking form of large, torn blisters. Further, the cellulose structure in M4 was also heavily altered. The structure of M5 resembled, to some extent M1 and M3; however, deep air channels instead of bubbles were observed. Cross sections of M6 were characterized by corrugated structure and numerous, very large, elongated air spaces due to their presence, the delamination was not as clearly visible as in case of M2. Finally, cross sections of M7 exhibited a greater degree of stratification, as compared to M2. Further, large air spaces, but no air bubbles, were observed between layers of this modified BC. In contrast to all of the modified BC pellicles, the control BC had no layers or blisters and was characterized by regular, compact structure, with overlapping layers of cellulose clearly visible. Thus, in the process of dehydration, the unmodified cellulose fibers flatten, undergoing irreversible deformation – this is the reason that dried, unmodified BC has significantly lower rehydration capacity.

In summary, SEM imaging revealed that crosslinking process results in two main changes in BC pellicle microstructure. First one, is evident stratification, created by the air spaces between successive layers of cellulose (e.g. M2, M7). Second one, is deformation caused by the presence of large, spherical or ellipsoidal air spaces (bubbles) (e.g. M4, M6). However, the most favorable SR (%) values (see **Figure S1**, **Tables S1-S9**) were obtained for BC samples where both types of BC microstructure alteration occurred simultaneously (M1, M3, M5). We conclude that the final microstructure of modified BC depends on the amount of gas produced and on the pressure generated within the sample during drying of fibers. It may be assumed that gradual release of smaller amounts of gas leads to BC delamination, while the rapid release of larger amounts of gas, leads to the formation of larger bubbles. In future studies, we plan to further examine the mechanism and specific gas released during crosslinking.

Simultaneously with SEM imaging, energy-dispersive X-ray spectroscopy (EDX) analysis was performed to determine the specific elemental composition of modified and control BC surfaces (**Figure S9**, **Table S10**). The measurements confirmed that carbon and oxygen were the main constituents of BC fibers, both in modified and unmodified samples. In the case of BC modified in the presence of CATs containing sodium salts (all the CATs except for CAT4), trace amount of sodium was also detected (0.2-0.5 w/w%). Somewhat surprisingly, nitrogen was not detected on the surface of BC modified with ammonium bicarbonate-based CATs (CATs 4-6). However, trace amounts of phosphorous were detected on M1, M3, and M5 samples, those that were modified with phosphorous-containing CATs (0.3-0.5 w/w%). It is worth noting, that in the case of the M7 sample (modified in the presence of sodium hypophosphite), the content of phosphorous was significantly higher (1.7 w/w%), as compared to the other aforementioned modifications.

### 3.3. Analysis of ATR-FTIR spectra of modified BC

Comparison of ATR-FTIR spectra of unmodified and modified BC indicated that all of the applied modification protocols had a major impact on BC chemical structure (**Figure 4A**, **Figure S10**). The primary difference as compared to non-treated BC was the appearance of a characteristic band at 1720 cm^−1^, indicating the presence of a carbonyl group (C=O, stretching) (61) – this can be considered a fingerprint of ester bonds and unreacted carboxylic groups of CA. The highest intensity of this band was observed following M5 modification, in which the combination of ammonium bicarbonate and disodium phosphate was used as catalyst, while for other the tested CATs the intensities were similar. Additionally, a second, characteristic and expected ester band appeared in all spectra of modified BC at approx. 1248 cm^−1^ (C-C-O asymmetric stretching) (62); however, its intensity did not vary between modification variants. Further, 2D-COS spectra (all modifications) also revealed that crosslinking process had a high impact on band intensities in the spectral region of 1200 cm^−1^ to 850 cm^−1^ (**Figure 4B**, **Figure 4C**). In this region, the spectra are primarily shaped by C-O-C stretching ether linkages and pyranose backbone rings and carbon-oxygen bonds (C-OH), indicating presence of primary alcohols (63). The reduction in intensity of bands in this region could be also a consequence of ester bond formation between carboxyl groups of citric acid (CA) and C-OH of anhydrous glucose units in BC fibrils. The presence of negative cross-peaks in the synchronous spectrum 1720 cm^−1^ vs. 950 cm^−1^ also suggests the possibility of a significant influence of modification process on the crystallinity of BC (64,65).

**Figure 4.**
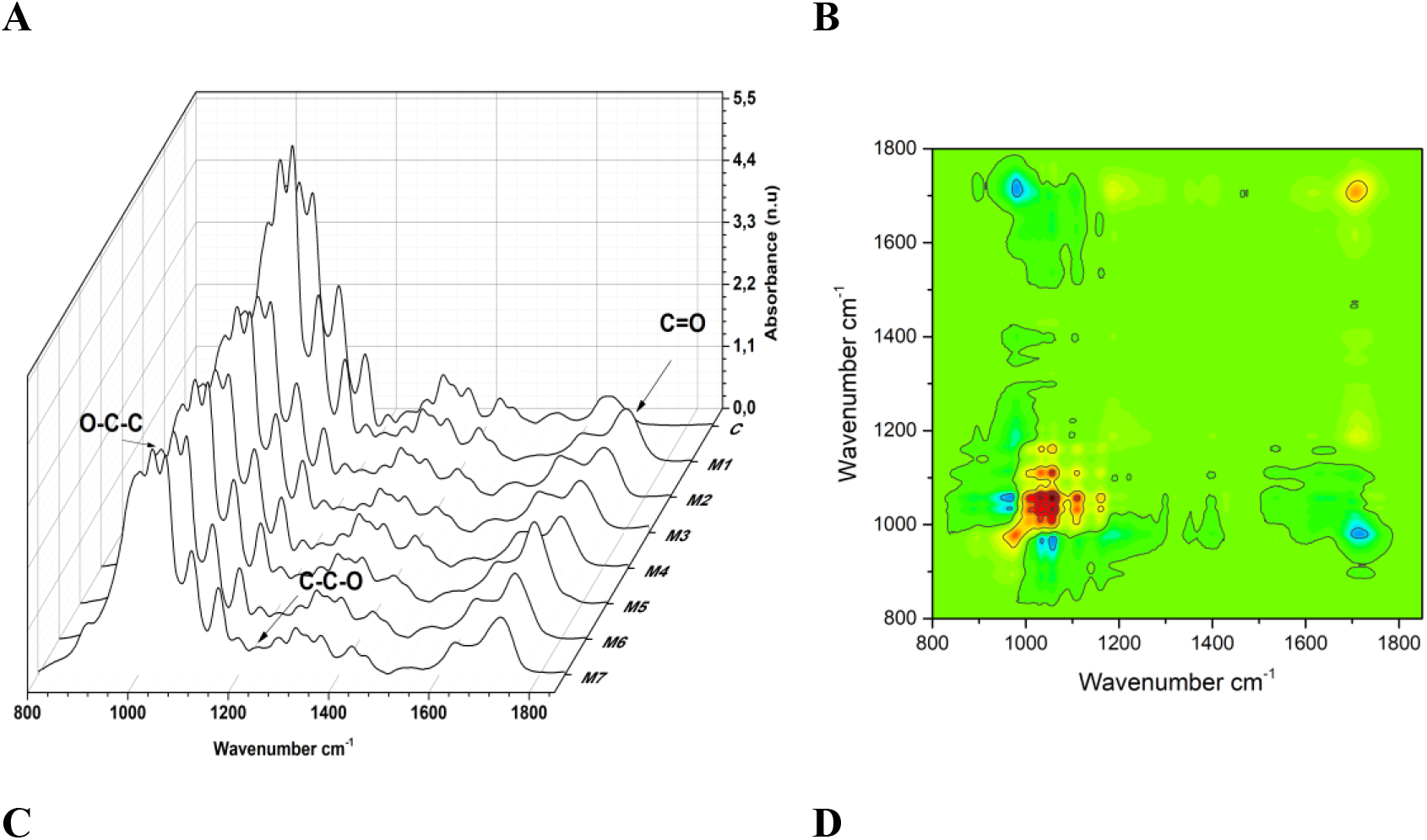
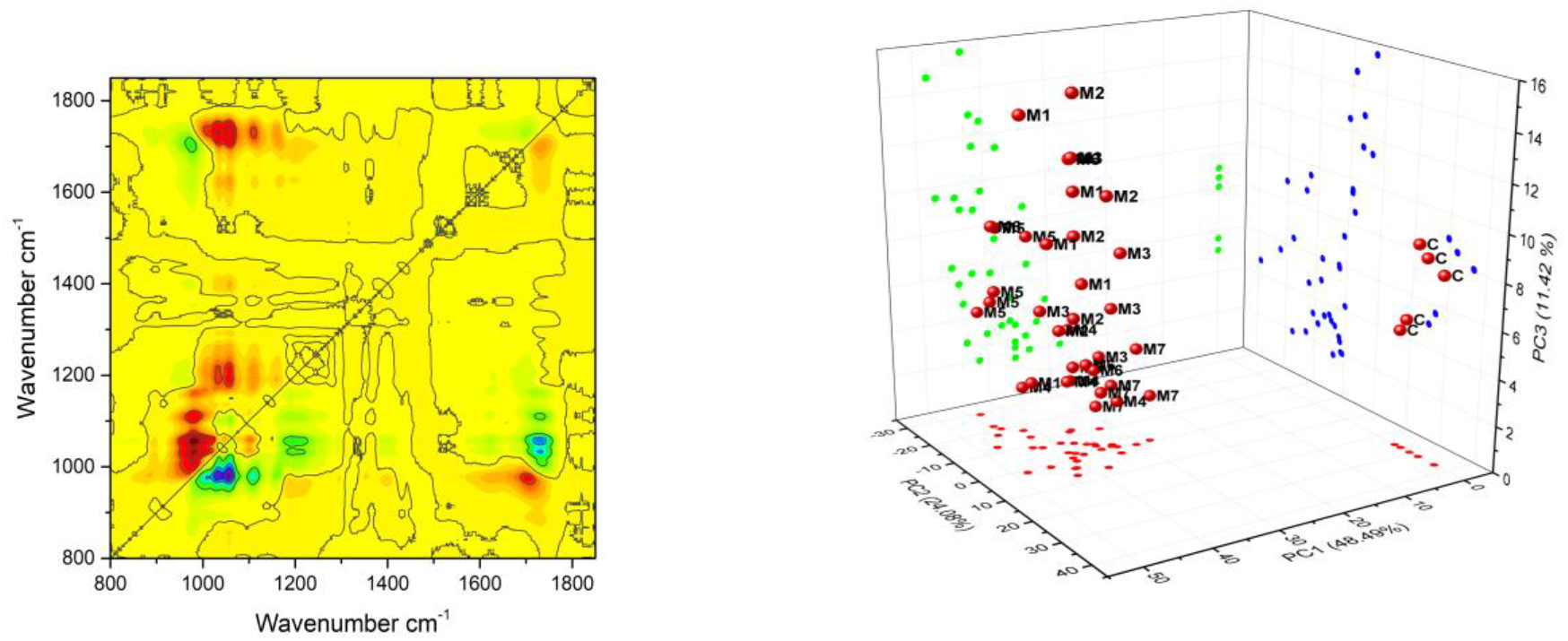
Result of ATR-FTIR analysis of crosslinked BC. **A)** 1D spectra region form 1800 cm^−1^ to 800 cm^−1^ of modified BC with different CATs; **B)** 2D – COS synchronous spectra, **C)** 2D – COS asynchronous spectra; **D)** Result of PCA with plotted PC1, PC2 and PC3 dimensions. C – control BC sample; M1-M7 – modified BC samples.

Finally, the preformed PCA of ATR-FTIR data confirmed successful modification of BC and showed also differences in the impact of the different types of modifications on the BC structure. From all extracted principal components, the first three account for approx. 80% of variance. In the PC1 (48.49%) dimension, the main contribution, accounting for over 80% of variance, was the result of changes at bands: 1320 cm^−1^, 1108 cm^−1^, 1277 cm^−1^, 1313 cm^−1^, 1427 cm^−1^, 1160 cm^−1^, 1248 cm^−1^, 1364 cm^−1^, 1720 cm^−1^. The changes in the first six bands influenced PC1 to the greatest degree, with almost equal contributions (above 10%). The PC2 (24.08%) dimension was determined primarily by set of bands in the spectral range from 1002 cm^−1^ to 895 cm^−1^, with the band at 980 cm^−1^ having the greatest influence on this dimension. The last dimension, PC3 (11.42%) was primarily determined by bands at 1002 cm^−1^, 1540 cm^−1^, 1640 cm^−1^, and 1720 cm^−1^. The 3D plot of all dimensions indicated that samples of control BC were a significant distance from samples of modified BC (**Figure 4D**). Further, samples of M1 and M2 were in similar region of space, in contrast to M5, M6, and M7. Finally, samples from M3 and M4 occupied the regions overlapping with previously listed modification variants.

### 3.4. Water-related properties and density of modified BC and efficiency of crosslinking reaction

The biomedical applications of BC as a wound dressing material depend primarily on its water-related properties, including water swelling and holding capacity, as well as water release rate. These properties depend, in turn, on pore size, pore volume, and surface area (66,67). The high liquid absorption capacity of BC is the primary driver for its application as a wound dressing material. Its excellent swelling capabilities not only help to absorb and sequestrate wound exudate, but also make dressing non-adherent to the wound bed. This latter feature is of paramount importance, not only with regard to easy, painless dressing removal, but also because it prevents the destruction of fresh, fragile granulation tissue (68,69). Further, the adsorption capacity of BC for liquids and small particles also makes it an appropriate material for impregnation with antimicrobials; therefore, it can act as a drug carrier with excellent wound healing potential (70). As previously explained, fully swollen or completely dried BC is significantly devoid of ability to absorb fluids, including wound exudate. However, we hypothesized that following crosslinking, dried BC would have preserved micro-fibrous structure and maintained high water absorption capacity.

The results presented in **Figure 5A**, (statistical analysis of these results is presented in **Table S11**) showed that after 60 min of incubation with water, SR (%) values for M1-M3 did not exceed twice the SR (%) value for the non-modified (control) BC. These samples were characterized by a slow, gradual increase in the SR (%) values over the entire period of 60 min and continued to absorb water up to 24 h (**Figure 5B**, statistical analysis of these results is presented in **Table S12**). After this time, SR (%) values for samples M1-M3 were more than 3 times higher compared to the control BC, for which no increase of absorption was observed between 60 min and 24 h incubation in water. The highest values of SR (%) were obtained for M3: approx. 2 times higher after 60 min and over 5 times higher after 24 h of incubation in water, as compared to control BC. For M4 after both 60 min and 24 h, the values of SR (%) for did not differ statistically from the control BC – after 24 h incubation in water only a slight increase in this parameter as was noticed. Interestingly, over the 60 min incubation, the M5 material exhibited a much higher increase in SR (%) than the control (by over 4 fold), as well as other modified BC samples. However, after further incubation in water no additionally water sorption was observed (the difference between 60 min and 24 h did not exceeded 1%), meaning that the maximum water absorption capacity of the M5 was reached after 60 min. Somewhat similarly, M6 samples were also characterized by a much higher value of the SR (%) after 60 min, as compared to the control (by over 3 fold). The swelling capacity of M6 after 60 min was lower only than for the M5, although the dynamics of water absorption was comparable between these two modifications. After 24 h, similarly to the M5, only a slight increase in SR (%) of M6 was found. The maximum SR (%) values of M6 were comparable with the M2 and M5. The sample modified with the reference catalyst (CAT7) after 60 min showed a degree of water absorption only slightly higher than the control. However, after 24 h, the SR (%) value for M7 sample was over 3 times higher as compared to the control. However, the maximum degree of water sorption for M7 was lower as compared to the M1 and M3 (approx. 1.5 and 1.7 times, respectively, with the differences being statistically significant), as well as to the M5 and M6 (approx. 1.3 times for both, but the differences were not statistically significant).

**Figure 5.**
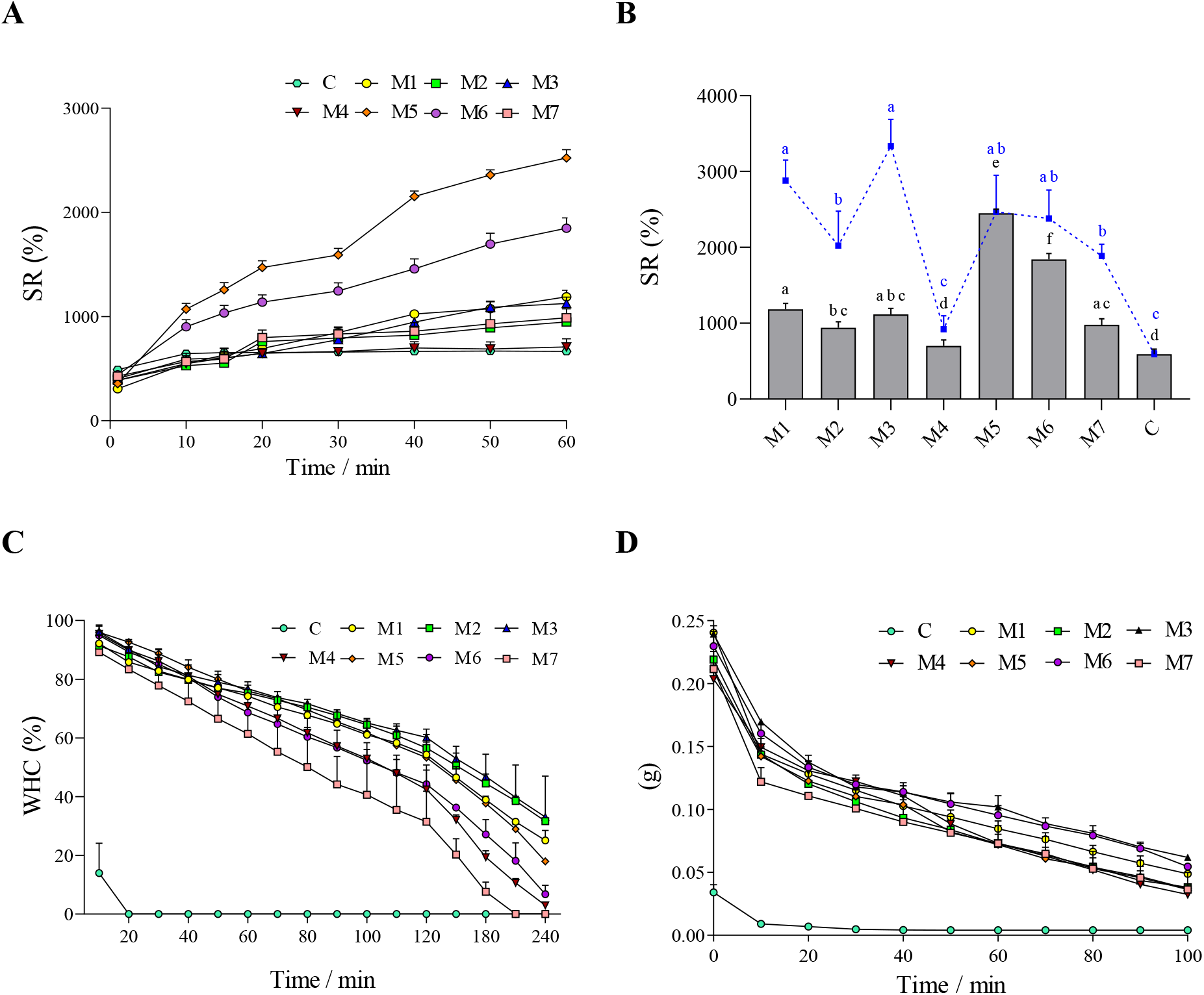
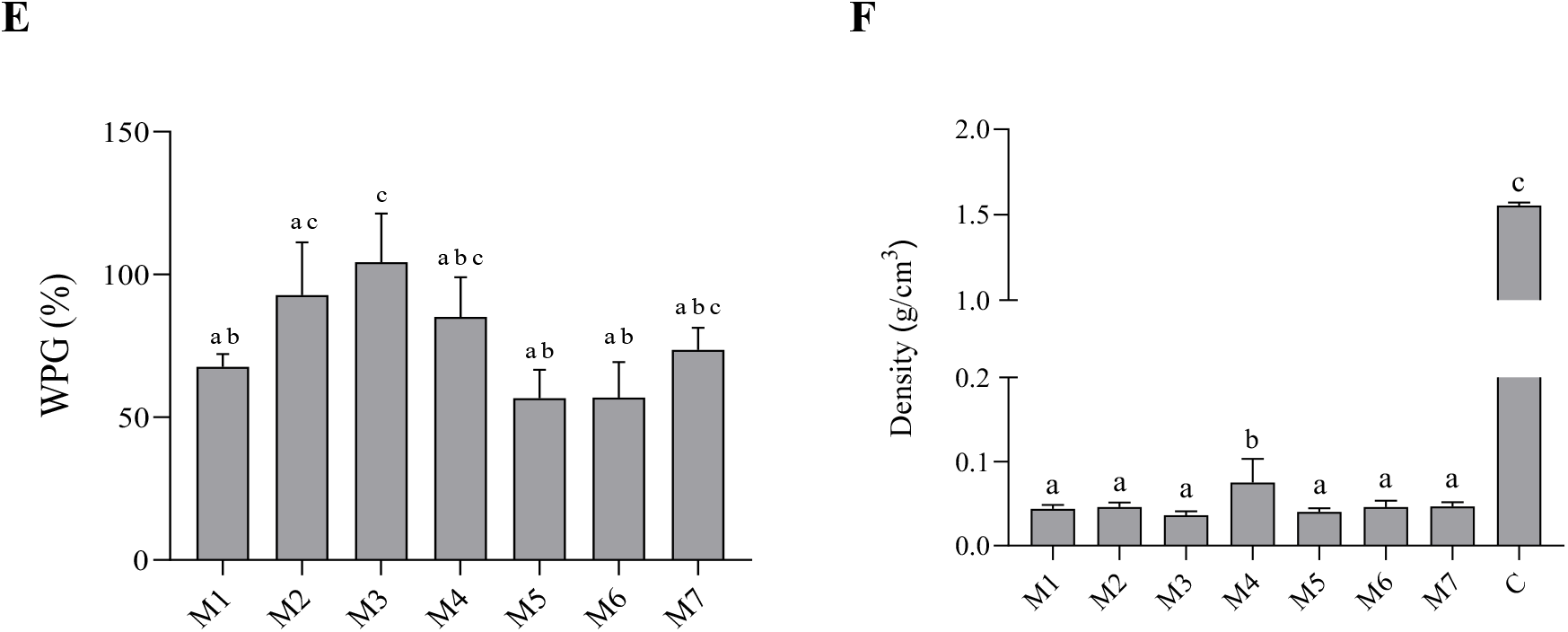
**A)** Swelling ratio (%) of BC samples over 60 min of incubation in water; **B)** Swelling ratio (%) of BC samples after 60 min of incubation in water (column) compared to swelling ratio (%) after 24 h of incubation in water (line); **C)** Water holding capacity (%) of BC samples over 60 min of incubation at 37°C; **D)** Weight loss of BC samples during centrifugation at 200 *g*; **E)** Efficiency of BC crosslinking reaction; **F)** Density of BC samples. Data are presented as mean± standard error of the mean (SEM); values with different letters are significantly different (p<0.05): a, b, c, d – statistically significant differences between the analyzed parameters. C – control BC sample; M1-M7 – modified BC samples.

As previously discussed, the BC chemical modifications studied here relied on citric acid (CA) crosslinking, facilitated by the presence of the different CATs, which released gas during the reaction with CA and as a result of thermal decomposition. This co-effect resulted in the formation of additional space within the cellulose matrix, enabling the high water sorption capacity of the modified BC samples after drying. At the same time the micro-porosity of the BC pellicles was unchanged, as compared to the unmodified control BC (**Figure S7**, **Figure S8B**).

The water molecules are trapped physically on the surface and inside the BC matrix consisting of reticulated fibrils (71). The more space is available between the BC fibrils, the more water can penetrate and adsorb onto the material. Further, the increase in the number and size of the pores results in an increase in surface area. The greater the surface area and the larger the pore size, the greater the capacity of BC to absorb water and swell (72,73).

Furthermore, in addition to an increase in swelling capacity, it was found that all of the modified BC samples demonstrated a significant increase in their capacity to hold water (**Figure 5C**, **Table S13**). This favorable water holding capacity of modified BC can also be ascribed to the differences in microstructure obtained as a result of the crosslinking proceeding with the different CATs. For unmodified BC, total water loss was observed after just 20 min of incubation at 37°C. Among modified samples, the quickest water loss was observed for the BC crosslinked with CAT7 (the WHC (%) reached 0 after 210 min). Samples M4 and M6 lost almost all absorbed water after 240 min, while BC modified with CATs 1-3 and CAT5 still held over 25% of their initial water volume after 240 min. Overall, the highest water retention (33%) after 240 min was observed for M3.

It has been demonstrated previously, that the amount of water lost from BC matrix to the outer environment depends on the arrangement of cellulose microfibrils (27). As demonstrated by Kaewnopparat et al. (74), only 10% of the water in BC behaves like free bulk water. Thus, most of the water molecules within the cellulose are, more or less tightly, bound to the cellulose. According to Ul-Islam et al. (22), BC samples with smaller pore sizes are able to retain water longer than those with larger pores. This effect is due to the closely arranged microfibrils binding water molecules more efficiently, with the stronger hydrogen bonding interactions (75,76).

Here, the gas released during the crosslinking reactions was responsible for the formation of numerous air cavities. It can be assumed that water filled up these spaces during WHC (%) analyses and exerted a pressure on the surrounding elastic cellulose fibrils, decreasing the pore sizes. This may explain why the modified samples M1-M3 in particular, absorbed water slowly, reaching maximum values of SR (%) after 24 h (**Figure 5B**). These assumptions are consistent with the observations from our previous work, in which we used vegetable oil to *in situ* modify BC (77). There, we hypothesized that during the process of BC development, cellulose compressed oil droplets and had an impact on the overall morphology of the environment, with oil droplets adapting a funnel-type form within the cellulose. However, at the same time, the hydrophobic oil pushed back on the BC matrix, consisting of >98% of water, thus compressing the fibrils and changing the overall morphology of the material.

Similarly, during centrifugation (200 *g*) the modified BC samples also held water significantly longer, as compared to the unmodified BC (**Figure 5D**, **Table S14**). The control sample released all absorbed water after 30 min of centrifugation, whereas all of the modified BC, regardless of the CAT used, retained more than 50% of the initial amount of water after 30 min of the process, and still retained some water after 100 min. However, in contrast to the experiment carried out at 37°C, in this experiment we did not observe any differences between BC samples modified using different CATs. Most likely, the forces affecting the material during centrifugation are much higher than those during evaporation at elevated temperature, and therefore the relatively small differences in the morphology of modified BC pellicles were not sufficient.

Finally, the efficiency of the crosslinking reactions in the presence of various CATs, expressed as weight percent gain (WPG (%)), was greatest for CAT3 (**Figure 5E**). This result was also correlated with these samples having the highest SR (%) value (**Figure 5B**). However, it should be noted that the M3 samples reached the maximum degree of water absorption after 24 h. Similar crosslinking efficiency was also found for reactions carried out in the presence of CAT2 and CAT4 (differences were statistically insignificant), but for the M4 modification, despite relatively similar efficiency, water absorption capacity was markedly lower (SR (%) values for M4 were comparable to unmodified control BC). Interestingly, the M5 and M6 modifications, which reached the maximum degree of water absorption after 60 min, had efficiency nearly 50% lower than CAT3 and with similar efficiency as with CAT1. Because the highest efficiency of the crosslinking reaction was observed in the presence of CAT3 (mixture of CAT1 and CAT2), it can be stated that the use of catalyst combinations can be more efficient with regard to the parameters discussed than application of single catalyst. However, the results obtained using CAT4, 5 and 6 showed these mixtures decrease the crosslinking efficiency, but still yield desired water sorption properties. Therefore, it cannot be stated that the effectiveness of the crosslinking reaction, measured as the weight gain of the material due to the binding of citric acid (CA) molecules, correlates with the values of SR (%) coefficient. However, this data does support our hypothesis that the primary role of the crosslinking agent is to stiffen the BC structure, while the increased ability to absorb water is essentially caused by the gasses released during thermal decomposition of CATs and the reaction of CA with CATs.

These considerations were supported by assessment of the densities of the modified BC pellicles (**Figure 5F**, **Table S15**). The M3 and M5 samples, previously characterized by the greatest ability to absorb water, had the lowest densities (over 30 times lower as compared to control). Interestingly, M3 and M5 yielded BCs that differed in the kinetics of water absorption and the effectiveness of the crosslinking reactions were not equal, but these differences were not reflected in their densities. In contrast, the M4 samples displayed the highest density (over 2 times higher as compared to other modifications) and absorbed the lowest volume of water. It is noteworthy, that the density of dried, unmodified BC was more than 15 times greater than M4, but no differences in water absorption were observed between these samples. Further, the efficiency of the crosslinking reaction in the presence of CAT4 did not differ statistically from reactions carried out with CAT3 and CAT5. However, these results can be explained the differences in the macro- and microstructures of these samples. As described above, CAT4 yielded high degree of BC delamination with lack of structural continuity – the air spaces were torn apart. Thus, these results confirm the key role of the formation of air spaces within BC layers, in improving the sorption properties of modified BC biomaterials, as well as the role of the crosslinking agent, which stiffens the BC structure and prevents it from collapsing during drying.

### 3.5. Assessment of cytotoxicity of modified BC

In line with use in wound dressings (surface contact with the body), we screened for cytotoxicity of modified BC pellicles *in vitro* by performing both extract and direct contact tests based on ISO 10993-5, as well as by assessing L929 fibroblast viability after culture on BC pellicles for 5 days using confocal microscopy. No evidence of cytotoxicity of any of the modified BC samples was observed (**Figure 6A**, **Figure S11**, **Table S17**) after 24 hours of culture of L929 fibroblasts with extracts (24 hours, complete growth media, 37°C). This indicated that no leachable toxic contaminants, such as residual citric acid or catalyst, were present, confirming that the purification protocol following modification was sufficient. Next, in the direct contact assay, where modified BC pellicle samples were placed on top of sub-confluent L929 cells in culture, we observed robust growth and normal cell morphology for control BC as well as M2, M3, M5, and M7 samples, while the cell density was somewhat reduced for the case of M1 and M6. However, for the case of M4 cell morphology was markedly altered and cell density was significantly lower (**Figure S12**, **Table S18**, **Table S19**). These microscopic observations were confirmed with the resazurin viability assay (**Figure 6A**). Overall, these results are consistent with the previous reports from Gyawali et al. (38), El Fawal et al. (39), that indicated that citric acid was safe and a nontoxic crosslinker for biomedical applications.

**Figure 6.**
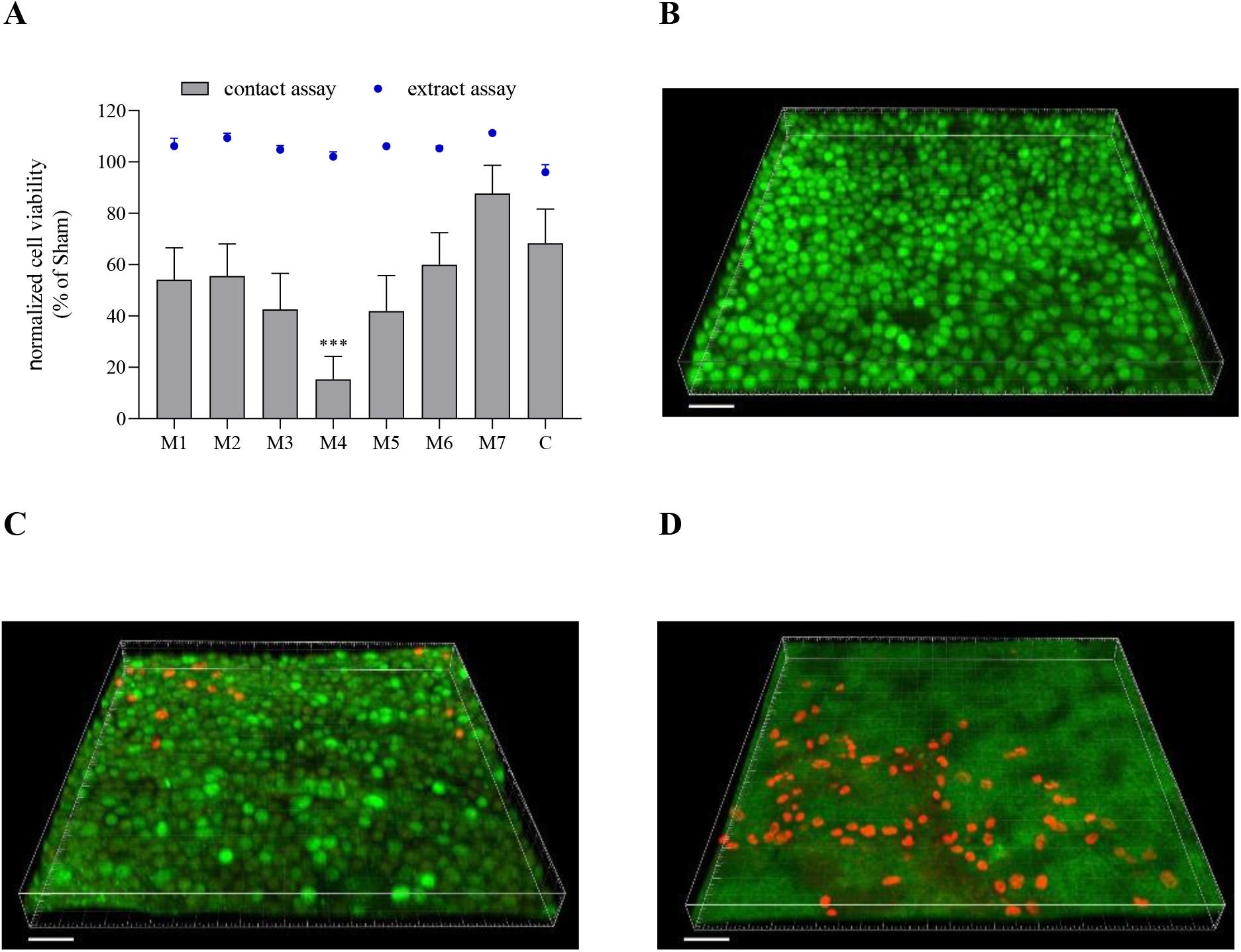
**A)** Viability data for all tested BC samples normalized to Sham (CellCrown insert for direct contact or complete growth medium for extract) obtained with resazurin viability assay; data are presented as mean ± standard error of the mean (SEM); the ‘***’ mark indicates statistical differences between modifications and control (p<0.001). **B-D**) Visualization of L929 fibroblasts cultured on control BC (B), M2 (C) and M4 (D) samples, respectively. Green color – nuclei in live fibroblasts (SYTO-9 staining, bright green) and exposed cell-free cellulose (dim green); red color – nuclei in dead fibroblasts (propidium iodide). Scale bar = 50 μm. C – control BC sample; M1-M7 – modified BC samples.

Finally, confocal microscopy (**Figure 6B-D**) of L929 cells cultured on modified BC pellicles for 5 days confirmed presence of live (green) fibroblasts on the surface of control BC, and showed predominantly live cells on M2 sample, with only a few dead cells. However, for the case of M5, we observed a considerable amount of dead (red) fibroblasts. Thus, this additional experiment aimed at evaluating cytotoxicity risk over a longer time period directly confirmed the results the extract and direct contact assays. Overall, the promising results obtained thus far motivate further studies of these materials as wound dressings in animals, aimed at assessing risk of sensitization and irritation.

To summarize, we tested and optimized a series of crosslinking reactions of BC using citric acid and various catalysts. Based on the collected results, we can single out use of a mixture of disodium phosphate and sodium bicarbonate (1:1 mass ratio, CAT3) as the most promising, yielding a SR (%) value of over 3300% after 24 hours. To our knowledge, the highest reported SR (%) value for modified BC was described by Figueiredo et al. (78). As reported by these authors, BC samples were impregnated with 2-aminoethyl methacrylate (AEM) with and without addition of N,N-methylenbis(acrylamide) (MBA), yielding SR (%) values reaching 6200%. However, in contrast to our modifications, the composites obtained by those authors were not dried before the swelling analyses. Additionally, while our highest SR (%) was lower as compared to the materials obtained by Figueiredo et al. (78), our crosslinking approach has some advantages. First of all, the use of an additional polymer, AEM, filled the porous BC structure completely, which may reduce gas exchange, a feature crucial for wound closure (79). Meanwhile, our M3-modified BC had preserved structure, including micro-porosity, and also contained numerous air cavities (see **Figure 3**, **Figure S2**, **Figure S7**). Another approach to obtain material with high sorption property described in the literature involves the use BC as a substrate for the production of hydrogel. As an example, Pandey et al. (80) obtained a hydrogel composed of BC and acrylamide that was characterized by SR (%) at neutral pH of 2500%, while Amin et al. (81) synthetized BC/acrylic acid-based hydrogel with a SR (%) value exceeding 5000%. However, to obtain these hydrogels, the BC had to be dissolved or ground into powder, and the synthesis process consisted of many stages. Additionally, it is important to consider the environmental health and safety aspects: acrylamide, acrylic acid and aminoethyl methacrylate are known to have toxic and irritant properties, limiting their applicability (82). Meanwhile, our M3 modification was obtained with use of simple and safe citric acid and disodium phosphate/sodium bicarbonate mixture. This fact cannot be neglected, especially considering the potential application of this BC material in wound dressings, where it will be in contact with fragile, exposed wound tissue.

To further assess the significance of our findings, we compared the swelling capacity of our M3-modified BC material, displaying the most promising features, to that of modern commercial dressings dedicated to highly exuding wounds. We found that the water swelling capacity of M3 (SR (%) over 3300%) was greater than that displayed by the market-leading modern dressings, e.g. polyacrylate fiber superabsorbent dressing (83) with SR (%) of 1400% or hydrofiber superabsorbent dressing (84) with SR (%) of 2600% (**Tables S20-S22**).

## 4. Conclusions

The use of citric acid crosslinker combined with the various inorganic CATs provides an important novel technique yielding significant improvement in BC water-related properties after drying. Our work and findings are of high impact for virtually all types of biomedical applications of BC. The compounds we tested as catalysts, with the exception of sodium hypophosphite, have never been used for crosslinking of BC and only disodium phosphate was previously used as bridging agent in general. We demonstrated that the formation of air cavities within the BC matrix played a key role in facilitating the absorption of large volumes of water. At the same time, our novel BC biomaterials released the absorbed water slowly, thanks to preserved micro-porous structure. These properties make the developed BC materials very promising for applications as wound dressings not only for highly-exuding, but also dry wounds (85,86). Moisture is an essential factor of wound healing process. However, excessive amount of exudate can macerate the wound bed, degrading peri-wound skin and growth factors, and increase the risk of inflammation (87). Our modified BC, in the dry state, may be applied as a “super-absorbent” wound dressing for exudate sequestration and maintenance of moisture levels appropriate for wound healing. Additionally, prior to drying, it is easy to envision loading the modified BC matrix with anti-inflammatory or antimicrobial drugs to be released as the dressing absorbs exudate. Alternately, in the case of dry wounds, the slow water release properties of our materials can be leveraged. For example, our modified BC biomaterial, chemisorbed with an isotonic, water-based fluid such as Ringer’s solution containing anti-inflammatory drugs, wound healing accelerators, or antimicrobial agents may restore proper moisture level and increase the rate of wound closure. Existing cellulose-based dressings saturated with such aqueous antimicrobial as polyhexanide (PHMB) or octenidine (OCT) are already used in the clinical setting (88,89).

We are aware that findings presented in this work are of *in vitro* character and require thorough testing in animal models before clinical trials may be performed. However, it should be emphasized that the application of cellulose-based dressings for purpose of exudate management is presently one of the most promising directions in chronic wound treatment (20,60). Bearing this in mind, the excellent results for materials using our novel modification, as compared to commercial dressings, indicate that the developed method has great potential to facilitate the design of innovative new dressings dedicated to chronic wound management.

## Supporting information

Supplementary data to this article

## Supporting Information

The results of BC crosslinking optimization, macro and micromorphological characteristics of modified BC, EDX spectra and BC elemental composition, PCA of ATR-FTIR spectra, assessment of cytotoxicity of modified BC – representative micrographs of L929 cells in direct/contact assay, comparison of swelling ability of modified BC with commercial superabsorbent dressings, statistical analysis of conducted experiments (PDF).

## Notes

The authors declare no competing financial interest.

## Acknowledgements

This work was supported by the National Centre for Research and Development in Poland (grant number LIDER/011/221/L-5/13/NCBR/2014).

## Abbreviations

BC: bacterial cellulose
CAT: catalyst
CA: citric acid
M1: modification with disodium phosphate as a catalyst (CAT)
M2: modification with sodium bicarbonate as a CAT
M3: modification with mixture of disodium phosphate and sodium bicarbonate in the ratio 1:1 as a CAT
M4: modification with ammonium bicarbonate as a CAT
M5: modification with disodium phosphate and ammonium bicarbonate in the ratio 1:1 as a CAT
M6: modification with sodium bicarbonate and ammonium bicarbonate in the ratio 1:1 as a CAT
M7: modification with sodium hypophosphite as a CAT
SHP: sodium hypophosphite
SR: swelling ratio
WHC: water holding capacity
WPG: weight percent gain.

