## Supplementary data to this article for "Superabsorbent crosslinked bacterial cellulose biomaterials for chronic wound dressings"

### Abbreviations

BC, bacterial cellulose; CAT, catalyst; CA, citric acid; M1, modification with disodium phosphate as a catalyst (CAT); M2, modification with sodium bicarbonate as a CAT; M3, modification with mixture of disodium phosphate and sodium bicarbonate in the ratio 1:1 as a CAT; M4, modification with ammonium bicarbonate as a CAT; M5, modification with disodium phosphate and ammonium bicarbonate in the ratio 1:1 as a CAT; M6, modification with sodium bicarbonate and ammonium bicarbonate in the ratio 1:1 as a CAT; M7, modification with sodium hypophosphite as a CAT; SHP, sodium hypophosphite; SR, swelling ratio; WHC, water holding capacity; WPG , weight percent gain.

### 1. Optimization of BC cross-linking reaction

The SR (%) values for BC samples modified using different CATs under the same reaction conditions displayed a comparable trend. The SR (%) results for the most effective M3 modification are presented in Figure S1, while the entire data set of the results from the optimization process tabulated in Tables S1 – S9.

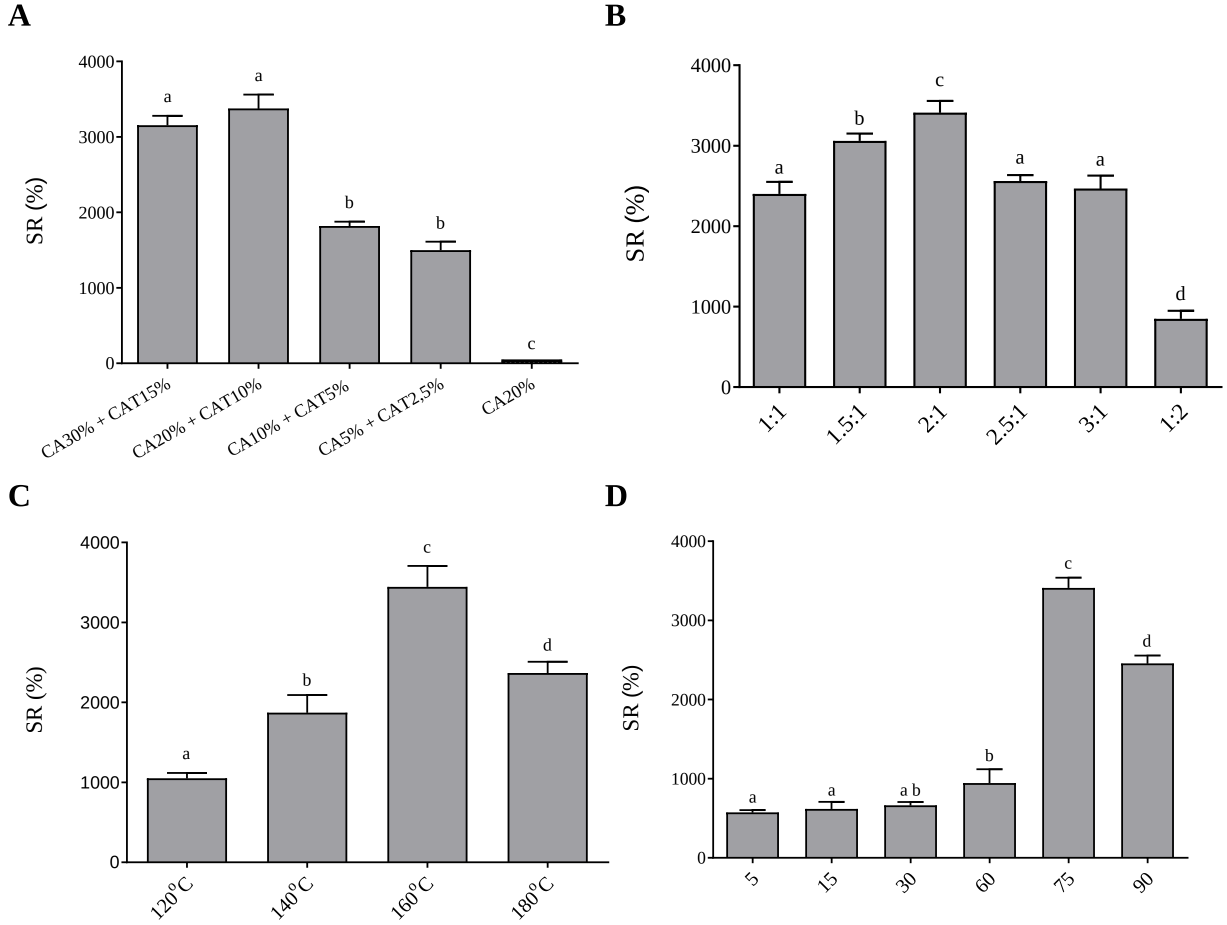


**Figure S1.** Swelling ratio (SR (%)) measured after 24 h, depending on the parameters used during optimization of BC crosslinking with CAT3 (disodium phosphate and sodium bicarbonate 1:1); **A** – different percentage of CA and CAT; **B** – different CA:CAT mass ratio; **C** – different reaction temperature; **D** – different reaction time.

Data are presented as mean ± standard error of the mean (SEM); values with different letters are significantly different ($p<0.05$): a, b, c, d – statistically significant differences between the analyzed parameters.

#### *1.1. Concentration of citric acid (CA) and catalysts (CAT) solutions*

Solutions consisting of 20% CA and 10% CAT, as well as 30% CA and 15% CAT – regardless CAT used – yielded BC pellicles with significantly higher values of SR (%), as compared to lower concentrations of both compounds (**Figure S1A**, **Table S1**, **Table S2**). Further, SR (%) did not differ significantly between 20% CA + 10% CAT and 30% CA + 15% CAT solutions. The highest SR (%) (3377.09 ± 184.43) was obtained for the M3. With decreasing CA and CAT concentration, below 20% CA and 10% CAT, a gradual decrease of SR (%) was observed. Thus, taking into account the economic aspect, for the further optimization stages the lower, but still highly effective, concentration of CA and CAT (CA20% + CAT10%) was chosen. These results are consistent with the results of Meftahi et al. (1), who reported the same concentration of CA and CAT (in their case, SHP) as the yielding the highest SR (%) values.

**Table S1.** Swelling ratio (SR (%)) of BC samples measured after 24 h of incubation in water depending on the percentage of citric acid (CA) and catalysts (CAT) solution.

| Type of modification | CA30% + CAT15% | CA20% + CAT10% | CA10% + CAT5% | CA5% + CAT2,5% |
|---|---|---|---|---|
| M1 | 2449.16 ± 156.45 | **2528.32 ± 192.08** | 1826.14 ± 100.02 | 1517.57 ± 62.19 |
| M2 | **2232.43 ± 112.60** | **2201.56 ± 50.66** | 1039.72 ± 46.89 | 933.58 ± 63.76 |
| M3 | 3154.53 ± 124.08 | **3377.09 ± 184.43** | 1818.91 ± 57.98 | 1499.65 ± 112.31 |
| M4 | **1084.74 ± 80.41** | **1082.45 ± 131.98** | 968.81 ± 68.64 | 966.31 ± 38.34 |
| M5 | 2292.05 ± 141.70 | **2324.66 ± 22.52** | 1786.63 ± 60.10 | 1186.33 ± 50.10 |
| M6 | 2247.87 ± 200.02 | **2343.81 ± 43.81** | 1556.29 ± 135.01 | 1176.79 ± 53.98 |
| M7 | 2016.87 ± 152.37 | **2033.25 ± 118.50** | 1245.54 ± 162.51 | 1049.19 ± 84.81 |

C – control BC sample; M1-M7 – modified BC samples.

**Table S2.** Statistical differences between SR (%) values presented in Table S1.

| | CA30% + CAT15% | CA20% + CAT10% | CA10% + CAT5% | CA5% + CAT2,5% | CA20% |
|---|---|---|---|---|---|
| CA30% + CAT15% | × | **ns** | **** | **** | **** |
| CA20% + CAT10% | ns | × | **** | **** | **** |
| CA10% + CAT5% | **** | **** | × | * | **** |
| CA5% + CAT2,5% | **** | **** | * | × | **** |
| CA20% | **** | **** | **** | **** | × |

* $p<0.05$, ** $p<0.01$, *** $p<0.001$, **** $p<0.0001$.

#### *1.2. Citric acid and catalysts solutions mass ratio*

BC samples modified using 20% CA + 10% CATs 1-5 and CAT7 reached the highest values of SR (%) at a CA:CAT mass ratio of 2:1 (**Figure S1B**, **Table S3**, **Table S4**). The highest values of SR (%) were obtained for M3 (3410.72 ± 146.97) and the SR (%) values were significantly higher, as compared to the other ratio variants. When CAT6 (mixture of sodium bicarbonate and ammonium bicarbonate) was used, the values of SR (%) obtained did not significantly differ between mass ratios 2:1 and 1.5:1. For the case of the M7, although the SR (%) reached the highest values at mass ratio 2:1, the further increase did not result in statistically significant differences (likewise between 1.5:1, 2.5:1 and 3:1). As a result, the 2:1 (CA:CAT) ratio was chosen for the further steps of the optimization. Additionally, our results are consistent with the CA:SHP ratio previously reported by Reddy & Yang (2).

**Table S3.** Swelling ratio (SR (%)) of BC samples measured after 24 h of incubation in water depending on the CA:CAT ratio.

| Type of modification | CA:CAT | | | | | |
|---|---|---|---|---|---|---|
| | 1:1 | 1.5:1 | 2:1 | 2.5:1 | 3:1 | 1:2 |

| | | | | | | |
|---|---|---|---|---|---|---|
| M1 | 2012.14 ± 109.11 | 2228.20 ± 62.50 | **2468.47 ± 34.69** | 2083.67 ± 62.41 | 2074.08 ± 69.13 | 743.71 ± 109.19 |
| M2 | 1887.34 ± 87.49 | 2010.88 ± 79.14 | **2178.67 ± 61.31** | 2036.58 ± 57.11 | 2073.20 ± 146.58 | 685.82 ± 85.04 |
| M3 | 2400.17 ± 147.02 | 3060.65 ± 89.58 | **3410.72 ± 146.97** | 2561.84 ± 73.33 | 2468.16 ± 160.54 | 849.62 ± 99.14 |
| M4 | 1001.73 ± 114.37 | 1031.81 ± 40.40 | **1110.47 ± 33.42** | 1101.47 ± 119.91 | 1069.14 ± 68.07 | 776.02 ± 105.40 |
| M5 | 2114.87 ± 102.96 | 2178.25 ± 22.70 | **2500.39 ± 47.87** | 2361.61 ± 201.80 | 2239.19 ± 10.88 | 839.18 ± 83.96 |
| M6 | 1944.51 ± 39.75 | **2403.25 ± 60.18** | **2399.71 ± 121.64** | 2172.80 ± 164.02 | 2090.97 ± 173.99 | 770.12 ± 48.43 |
| M7 | 1888.74 ± 88.38 | 1945.37 ± 179.11 | **2028.26 ± 57.11** | 2020.59 ± 50.41 | 1985.56 ± 86.06 | 866.32 ± 108.60 |

C – control BC sample; M1-M7 – modified BC samples.

**Table S4.** Statistical differences between SR (%) values presented in Table S3.

| | 1:1 | 1.5:1 | 2:1 | 2.5:1 | 3:1 | 1:2 |
|---|---|---|---|---|---|---|
| 1:1 | × | *** | **** | ns | ns | **** |
| 1.5:1 | *** | × | * | ** | *** | **** |
| 2:1 | **** | * | × | **** | **** | **** |
| 2.5:1 | ns | ** | **** | × | ns | **** |
| 3:1 | ns | *** | **** | ns | × | **** |
| 1:2 | **** | **** | **** | **** | **** | × |

* p<0.05, ** p<0.01, *** p<0.001, **** p<0.0001.

#### *1.3. Reaction temperature*

Regardless of the CAT used (**Figure S1C**, **Table S5**, **Table S6**) the highest SR (%) was obtained at a temperature of 160°C. Overall, increasing the temperature up to 160°C correlated with increase of SR (%) in all BC samples, but when the temperature exceeded 160°C, the SR (%) decreased.

**Table S5.** Swelling ratio (SR (%)) of BC samples measured after 24 h of incubation in water depending on the reaction temperature.

| Type of modification | Reaction temperature (°C) | | | |
|---|---|---|---|---|
| | 120 | 140 | 160 | 180 |
| M1 | 844.87 ± 141.14 | 1605.45 ± 230.53 | **2479.11 ± 97.01** | 1834.64 ± 78.82 |

| | | | | |
|---|---|---|---|---|
| M2 | 911.85 ± 81.53 | 1235.85 ± 75.58 | **2193.29 ± 371.01** | 1832.06 ± 133.83 |
| M3 | 1051.64 ± 65.72 | 1871.51 ± 221.02 | **3444.81 ± 262.25** | 2367.43 ± 139.72 |
| M4 | 680.51 ± 102.15 | 857.58 ± 75.31 | **1101.29 ± 46.17** | 961.62 ± 35.76 |
| M5 | 1087.18 ± 150.58 | 1491.20 ± 49.00 | **2223.27 ± 117.86** | 2009.50 ± 93.58 |
| M6 | 918.38 ± 95.86 | 1722.73 ± 134.25 | **2265.81 ± 363.48** | 1883.16 ± 89.70 |
| M7 | 707.86 ± 92.64 | 802.36 ± 88.81 | **2019.84 ± 75.91** | 1315.95 ± 187.29 |

C – control BC sample; M1-M7 – modified BC samples.

**Table S6.** Statistical differences between SR (%) values presented in Table S5.

| | 120 | 140 | 160 | 180 |
|---|---|---|---|---|
| 120 | × | ** | **** | *** |
| 140 | ** | × | **** | * |
| 160 | **** | **** | × | *** |
| 180 | *** | * | *** | × |

* p<0.05, ** p<0.01, *** p<0.001, **** p<0.0001.

#### *1.4. Duration of crosslinking reaction*

Once again, regardless of the CAT used, the highest SR (%) values were obtained after 75 min of crosslinking reaction (**Figure S1D**, **Table S7**, **Table S8**). However, it is important to note that the duration of crosslinking must be adjusted to the weight of the sample – the heavier it is, the longer time of crosslinking is required (data not shown). Here, the BC samples had an average weight of 4.5 g. Compared to the work of Meftahi et al. (1), where BC samples were crosslinked at 160°C for just 5 min, the optimal crosslinking duration determined here was significantly longer, however the precise size and weight of BC samples used in that work were not clearly specified.

**Table S7.** Swelling ratio (SR (%)) of BC samples measured after 24 h of incubation in water depending on the time of reaction carried out at 160°C.

| Type of modification | Reaction time (min) | | | | | |
|---|---|---|---|---|---|---|
| | 5 | 15 | 30 | 60 | 75 | 90 |

| | | | | | | |
|---|---|---|---|---|---|---|
| M1 | 529.57 ± 28.20 | 632.38 ± 51.50 | 668.04 ± 50.72 | 928.64 ± 88.97 | **2417.22 ± 122.38** | 1621.78 ± 35.97 |
| M2 | 629.98 ± 42.83 | 751.51 ± 30.88 | 816.66 ± 36.72 | 937.05 ± 154.54 | **2037.51 ± 74.38** | 1146.15 ± 51.36 |
| M3 | 573.89 v 28.77 | 618.04 ± 89.14 | 662.94 ± 42.53 | 944.21 ± 176.28 | **3410.32 ± 130.04** | 2456.54 ± 98.43 |
| M4 | 523.39 ± 44.48 | 602.04 ± 17.57 | 659.34 ± 40.00 | 712.83 ± 62.14 | **989.52 ± 58.44** | 915.05 ± 71.75 |
| M5 | 568.39 ± 24.09 | 633.18 ± 58.49 | 628.67 ± 14.74 | 918.83 ± 128.36 | **2229.65 ± 106.17** | 1711.16 ± 137.43 |
| M6 | 603.73 ± 46.82 | 611.33 ± 21.59 | 562.30 ± 73.09 | 1095.10 ± 12.13 | **2340.193 ± 196.60** | 1974.74 ± 101.48 |
| M7 | 542.68 ± 48.27 | 630.32 ± 53.38 | 632.16 ± 98.74 | 879.46 ± 57.53 | **2018.44 ± 166.23** | 1508.25 ± 173.19 |

C – control BC sample; M1-M7 – modified BC samples.

**Table S8.** Statistical differences between SR (%) values presented in Table S7.

| | 5 | 15 | 30 | 60 | 75 | 90 |
|---|---|---|---|---|---|---|
| 5 | × | ns | **ns** | * | **** | **** |
| 15 | ns | × | **ns** | * | **** | **** |
| 30 | ns | ns | × | ns | **** | **** |
| 60 | * | * | **ns** | × | **** | **** |
| 75 | **** | **** | **** | **** | × | **** |
| 90 | **** | **** | **** | **** | **** | × |

* p<0.05, ** p<0.01, *** p<0.001, **** p<0.0001.

**Table S9.** Adjusted p-values for each variant of optimization.

| | * | ** | *** | **** |
|---|---|---|---|---|
| **Percentage of CA and CAT solutions** | 0.0412 | - | - | < 0.0001 |
| **CA and CAT solutions mass ratio** | 0.0444 | 0.0039 | 0.0003 – 0.009 | < 0.0001 |
| **Reaction temperature** | 0.0481 | 0.0031 | 0.0001 – 0.0005 | < 0.0001 |
| **Reaction time** | 0.0111 – 0.0261 | - | - | < 0.0001 |

### 2. Macro and micromorphological characteristics of modified BC

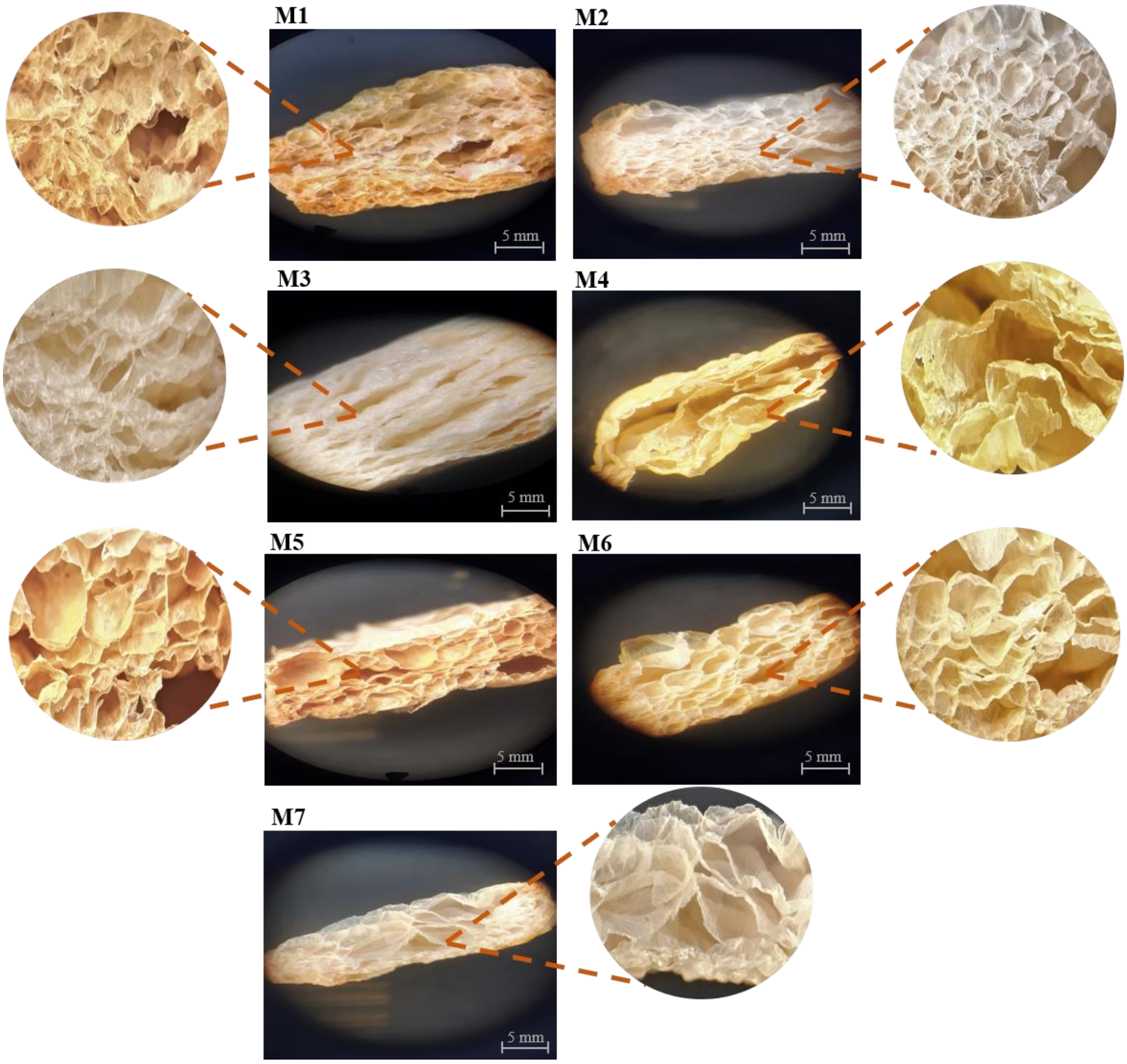


**Figure S2.** Cross sections of modified BC samples (magnification 5x and 25x, pictures in the center and on the edges of the figure, respectively) Pictures were taken using stereoscopic microscope (MST, Zeiss, Oberkochen, Germany). C – control BC sample; M1-M7 – modified BC samples.

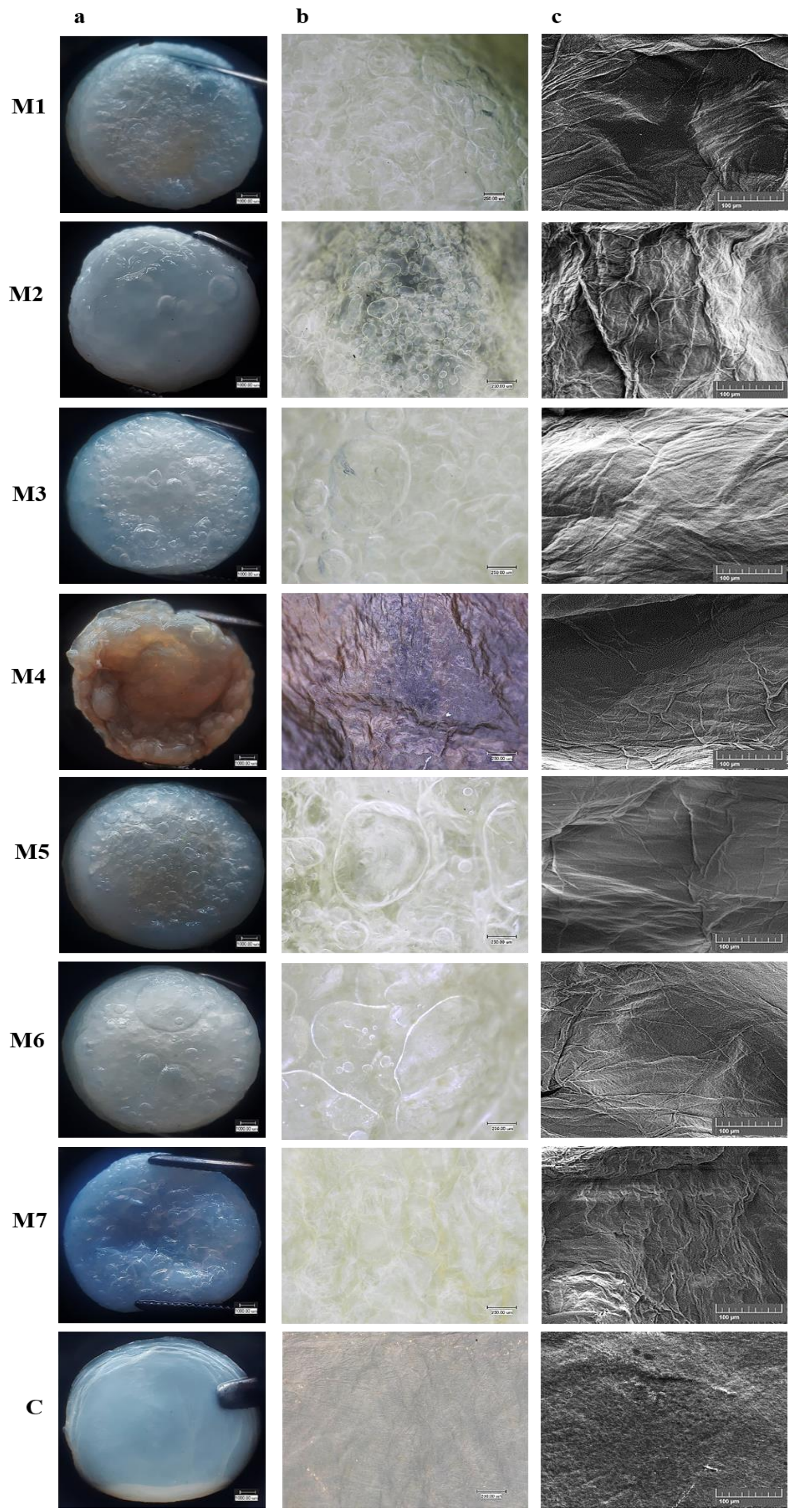


**Figure S3.** The surface of modified and unmodified (control) BC samples. a, b – magnification 20x and 150x, respectively (stereoscopic microscope, MST, Zeiss, Oberkochen, Germany); c –

magnification 300x (SEM, Auriga 60, Zeiss, Oberkochen, Germany). C – control BC sample; M1-M7 – modified BC samples.

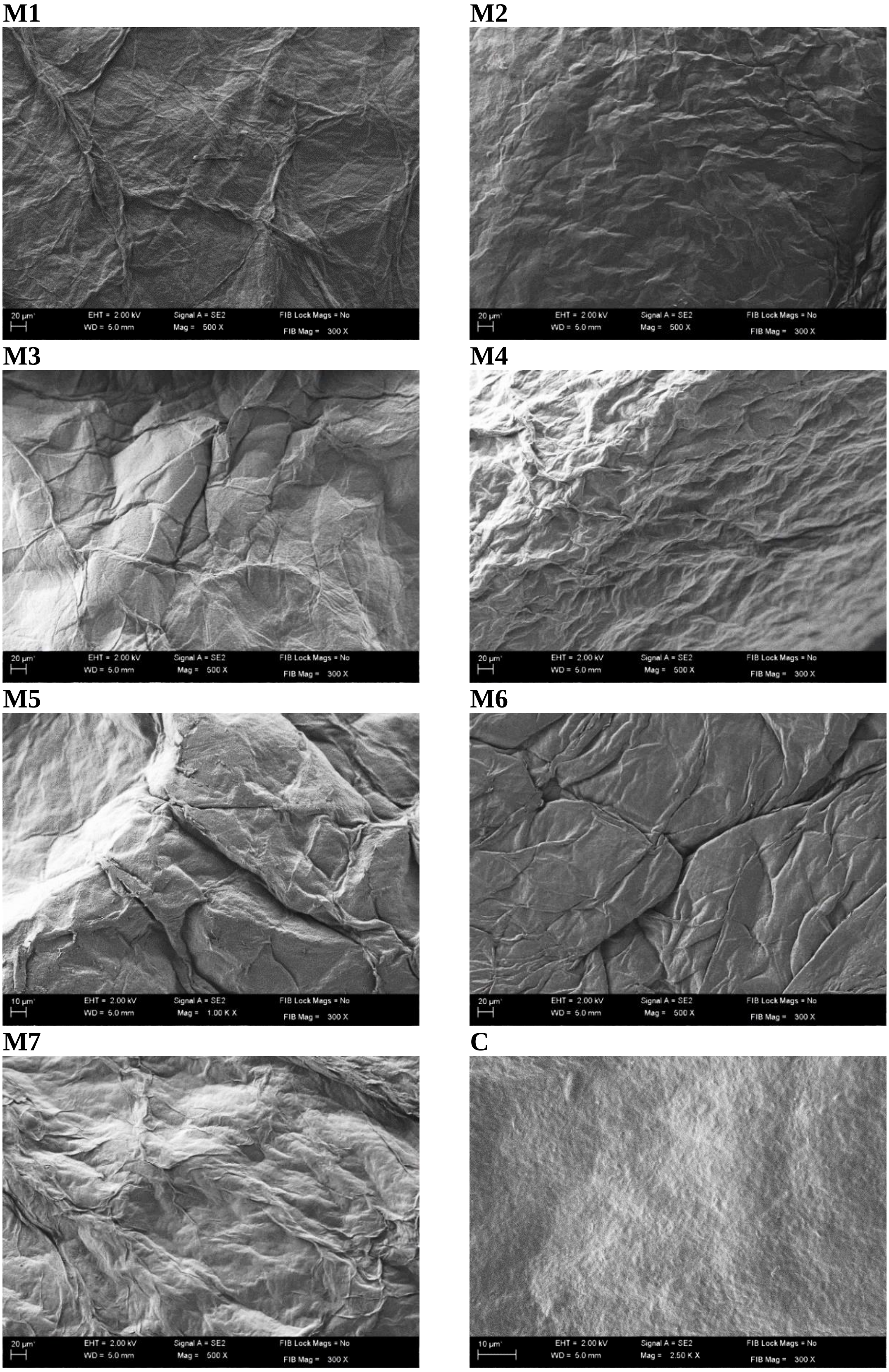

**Figure S4.** The surface of modified and unmodified (control) BC samples (magnification 500x, SEM, Auriga 60, Zeiss, Oberkochen, Germany). C – control BC sample; M1-M7 – modified BC samples.

**M1**

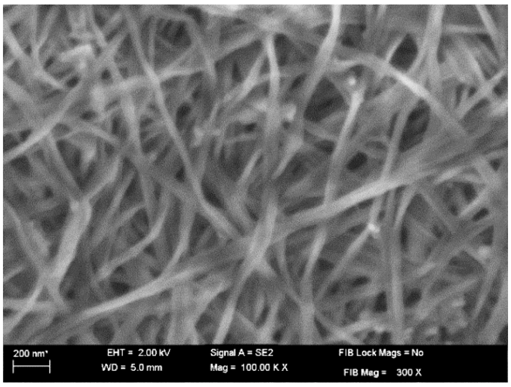


**M2**

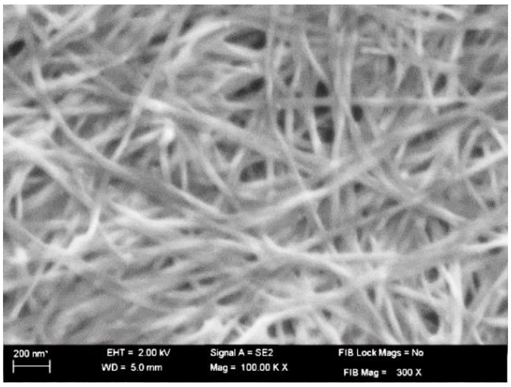


**M3**

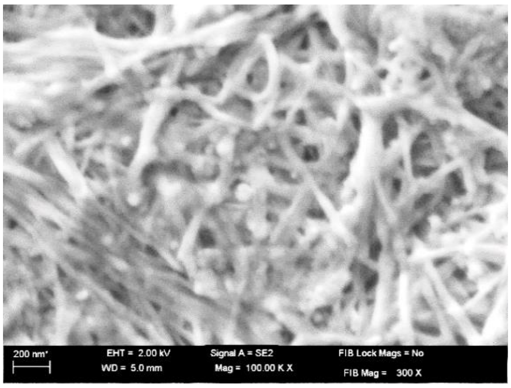


**M4**

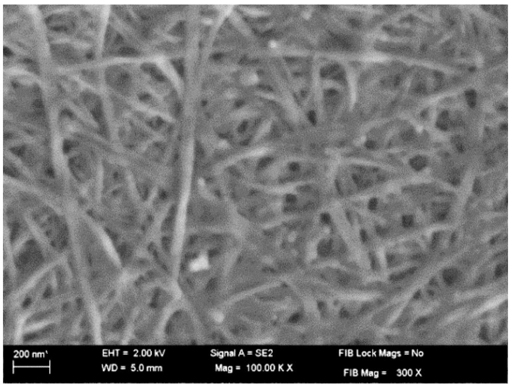


**M5**

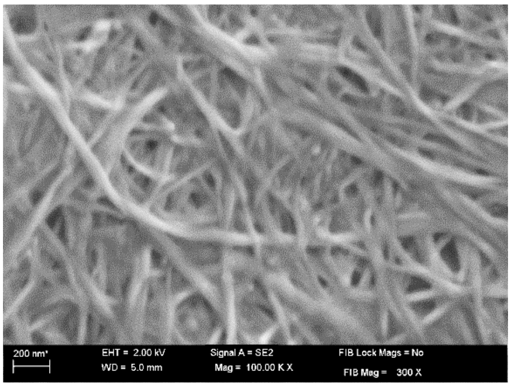


**M6**

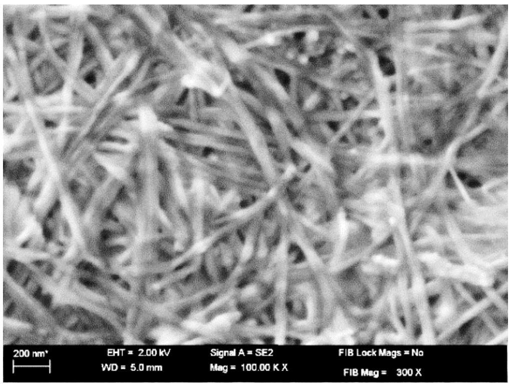


**M7**

**C**

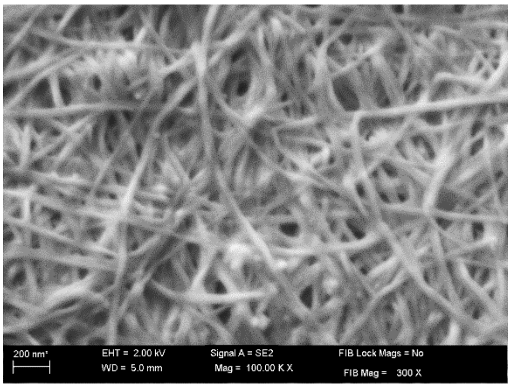


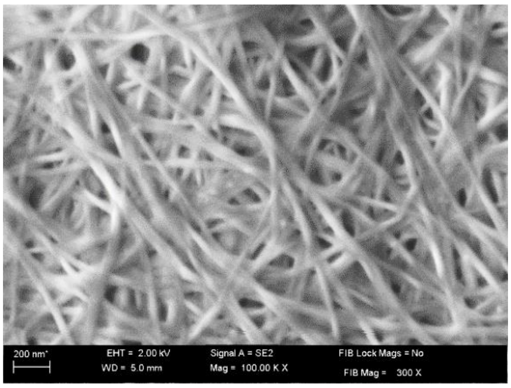


**Figure S5.** The microstructure of the surface of modified and unmodified (control) BC samples (magnification 10 000x, SEM, Auriga 60, Zeiss, Oberkochen, Germany). C – control BC sample; M1-M7 – modified BC samples.

**M1** **M2**

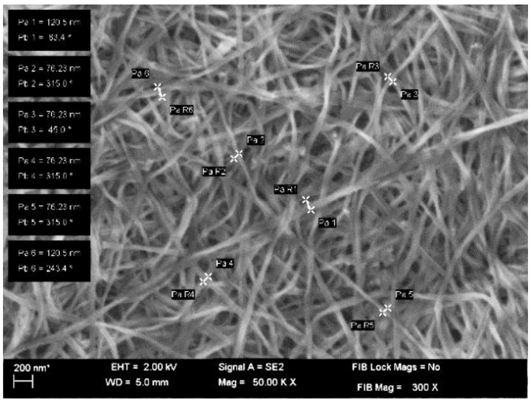


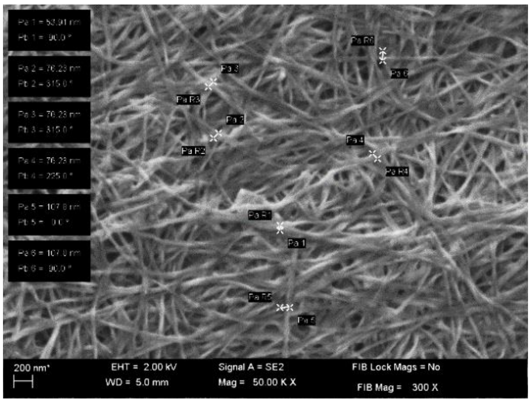


**M3** **M4**

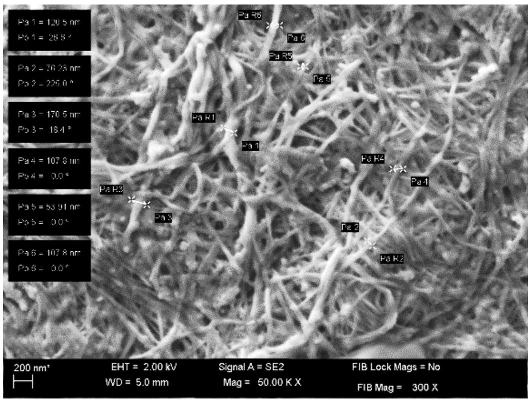


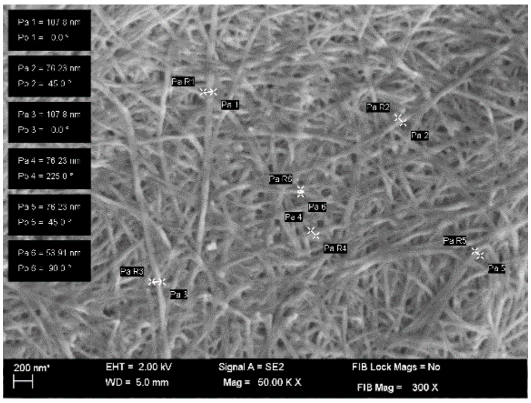


**M5** **M6**

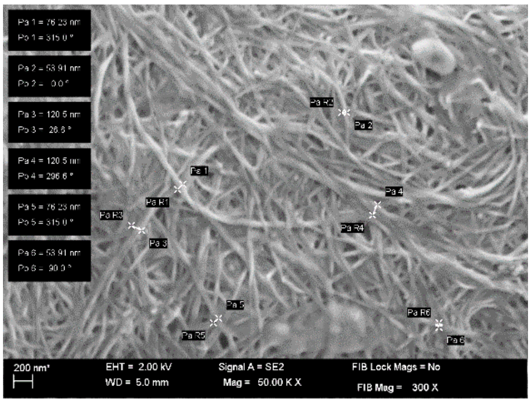


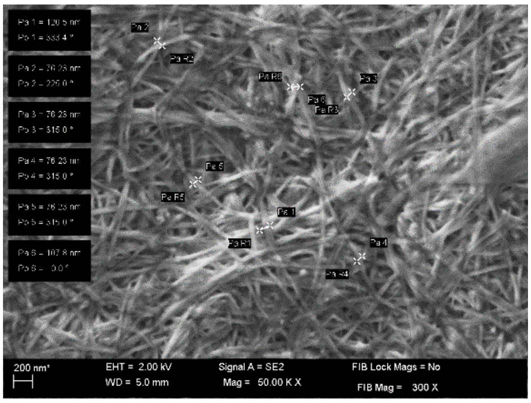


**M7** **C**

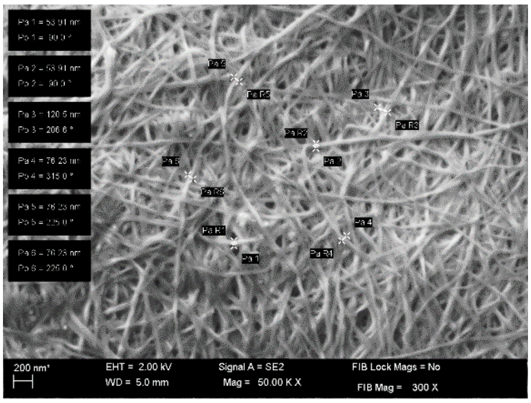


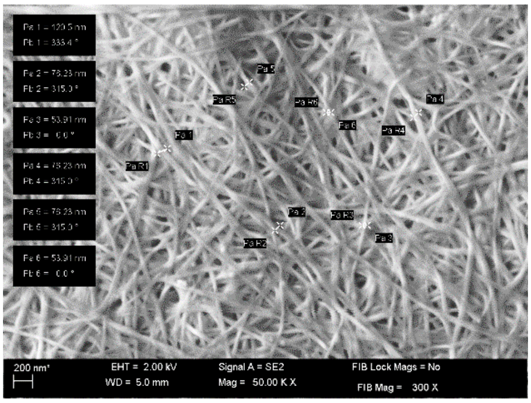


**Figure S6.** SEM images of modified and unmodified (control) BC samples with microfibrils diameters marked (magnification 5 000x, SEM, Auriga 60, Zeiss, Oberkochen, Germany). C – control BC sample; M1-M7 – modified BC samples.

**M1** **M2**

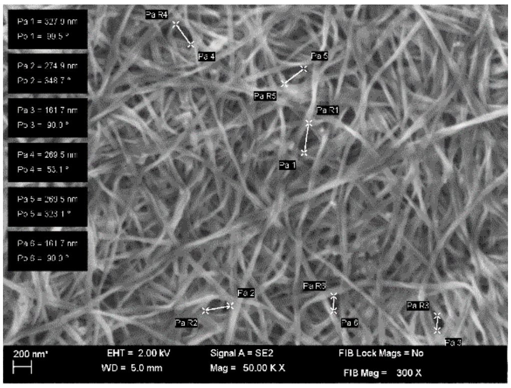


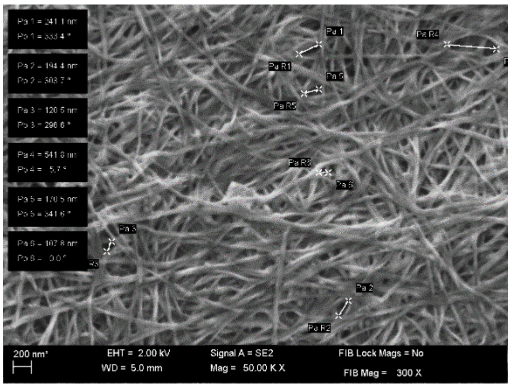


**M3** **M4**

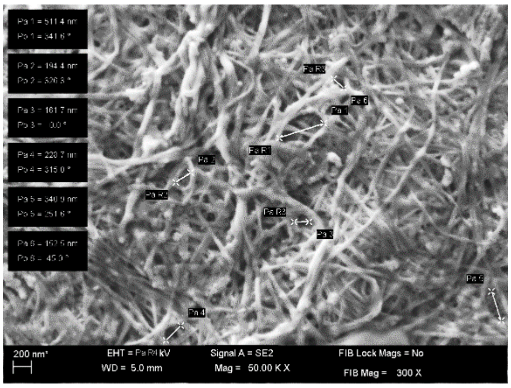


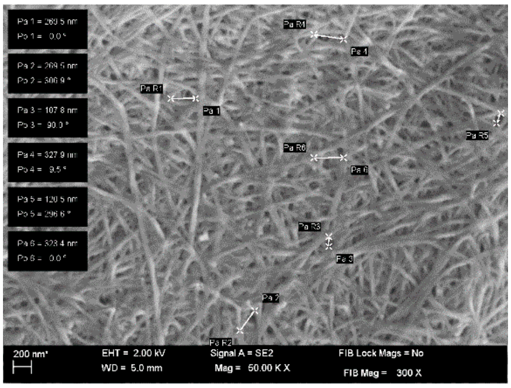


**M5** **M6**

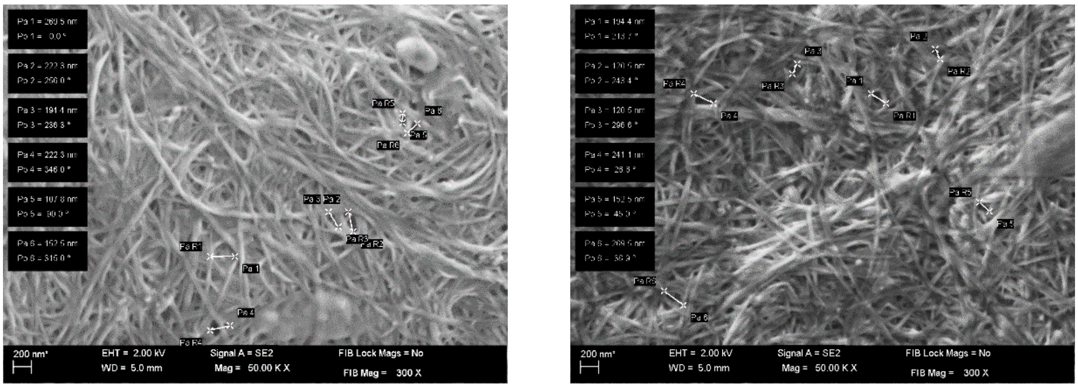


**M7**

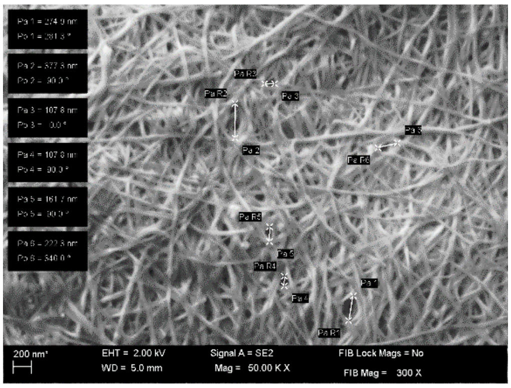


**C**

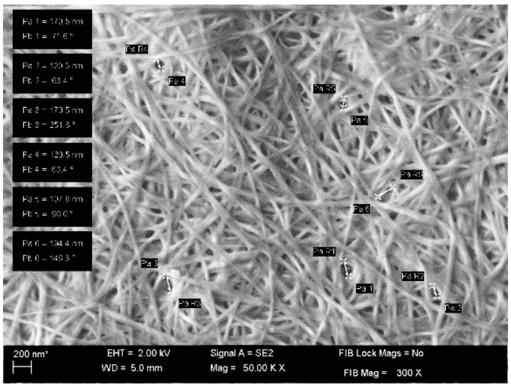


**Figure S7.** SEM images of modified and unmodified (control) BC samples with pore diameters marked (magnification 5 000x, SEM, Auriga 60, Zeiss, Oberkochen, Germany). C – control BC sample; M1-M7 – modified BC samples.

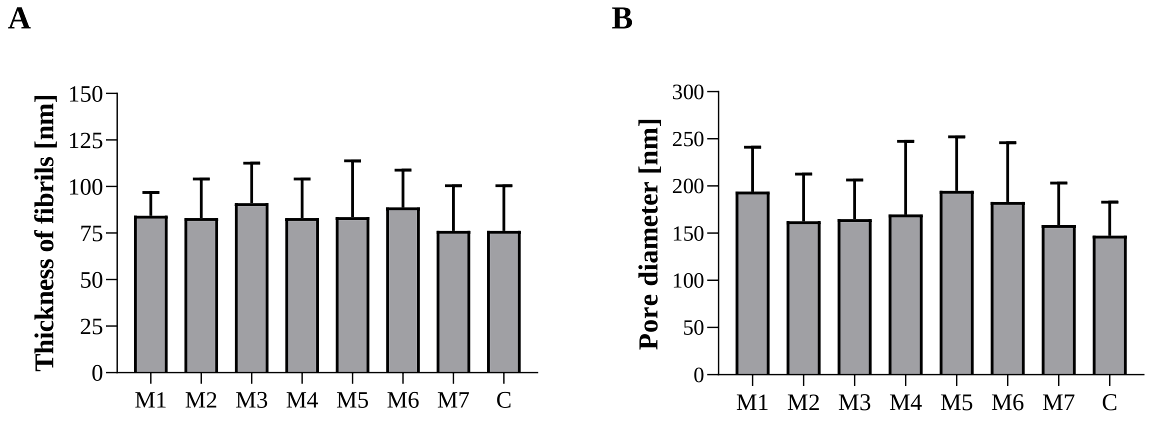


**Figure S8**. The thickness and pore diameter measured on the surface of modified and unmodified (control) BC samples. The BC microfibril diameters were analyzed using software integrated with Auriga 60 SEM (Zeiss, Oberkochen, Germany), while pore size was analyzed using ImageJ software (NIH). Data are presented as mean ± standard error of the mean (SEM). There were no statistically significant differences between the samples. C – control BC sample; M1-M7 – modified BC samples.

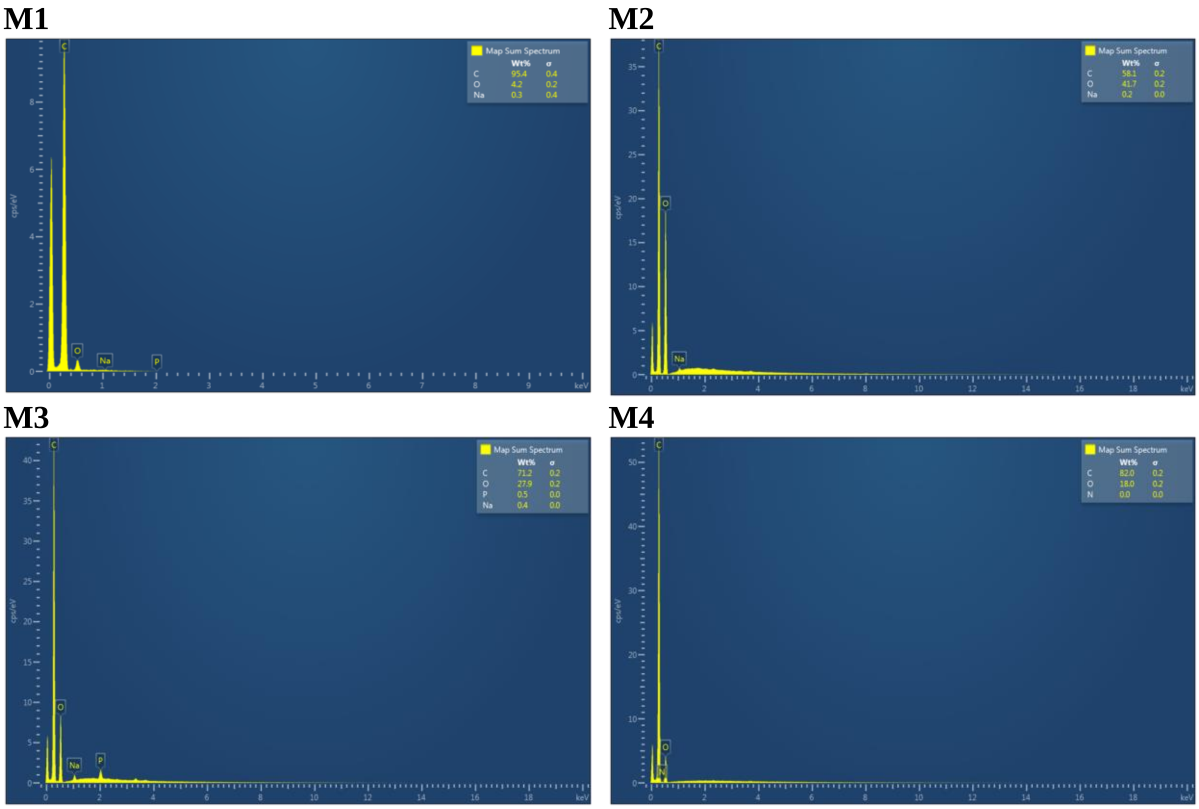

**M5** **M6** **M7** **C**

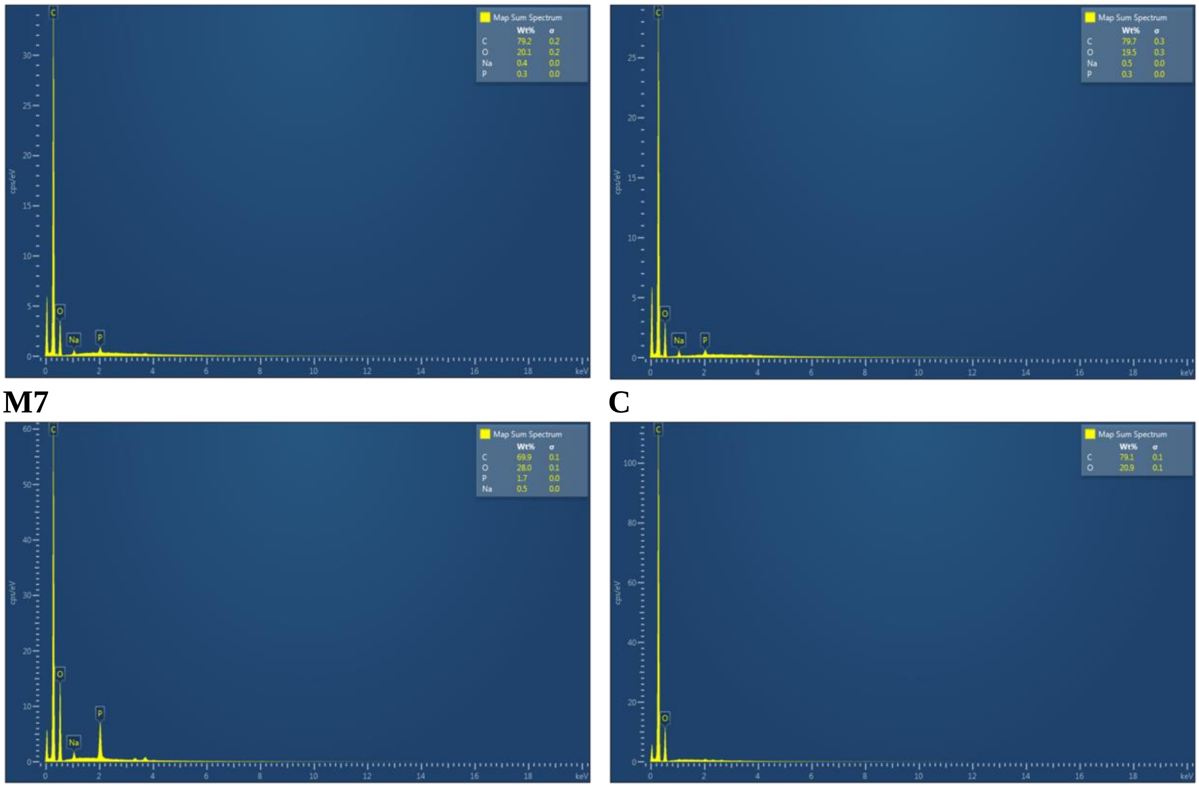

**Figure S9**. EDX spectra of modified and unmodified (control) BC samples (AZtec system, Oxford Instruments, Abingdon, United Kingdom). C – control BC sample; M1-M7 – modified BC samples.

**Table S10.** Results of EDX analysis.

| | | M1 | M2 | M3 | M4 | M5 | M6 | M7 | C |
|---|---|---|---|---|---|---|---|---|---|
| $W_t$% | C | 73.7 | 58.1 | 71.2 | 82.0 | 79.2 | 79.7 | 69.9 | 79.1 |
| | O | 25.4 | 41.7 | 27.9 | 18.0 | 20.1 | 19.5 | 28.0 | 20.9 |
| | Na | 0.4 | 0.2 | 0.4 | - | 0.4 | 0.5 | 0.5 | - |
| | P | 0.5 | - | 0.5 | - | 0.3 | 0.3 | 1.7 | - |
| | N | - | - | - | - | - | - | - | - |

$W_t$% - Elemental composition (%); C – control BC sample; M1-M7 – modified BC samples.

### 3. Principle Component Analysis of ATR-FTIR spectra

**A**

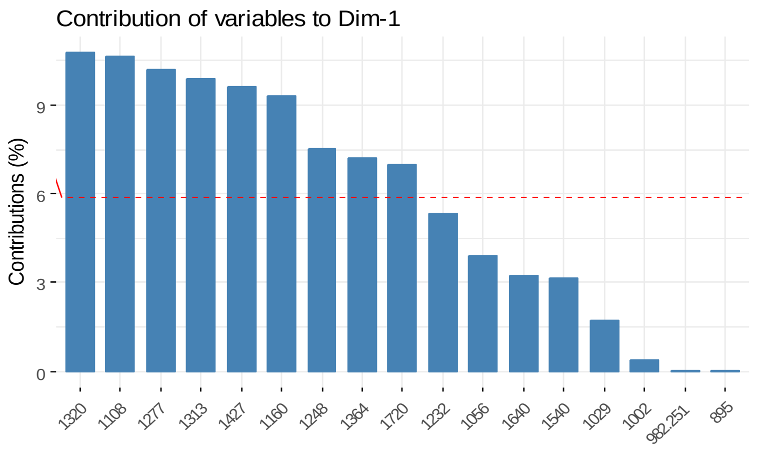


**B**

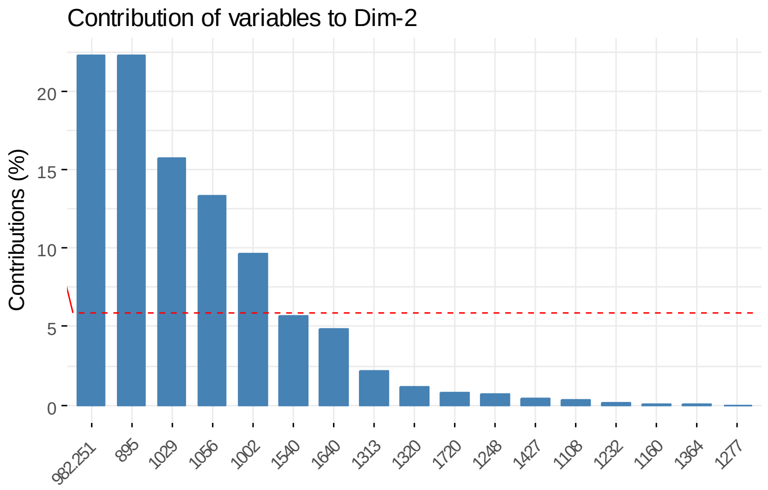


**C**

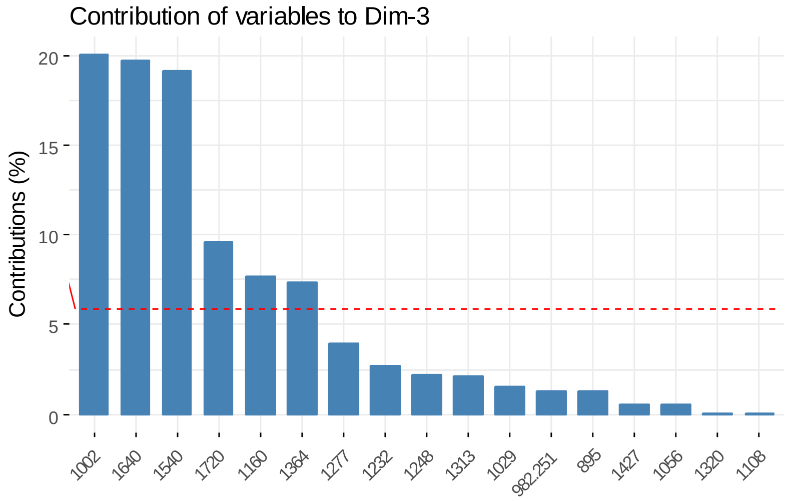


**Figure S10.** Contribution of variables according to the results from ATR-FTIR analyses.

### 4. Water-related properties and density of modified BC and efficiency of crosslinking reaction

**Table S11.** Statistical differences between SR (%) values obtained after 60 min of incubation of BC samples in water.

| | M1 | M2 | M3 | M4 | M5 | M6 | M7 | C |
|---|---|---|---|---|---|---|---|---|
| M1 | × | ns | ns | *** | **** | **** | ns | **** |
| M2 | ns | × | ns | ns | **** | **** | ns | * |
| M3 | ns | ns | × | ** | **** | **** | ns | *** |
| M4 | *** | ns | ** | × | **** | **** | ns | ns |
| M5 | **** | **** | **** | **** | × | ns | *** | **** |
| M6 | **** | **** | **** | **** | ns | × | ns | **** |
| M7 | ns | ns | ns | ns | **** | **** | × | ** |
| C | **** | * | *** | ns | **** | **** | ** | × |

C – control BC sample; M1-M7 – modified BC samples; * $p<0.05$, ** $p<0.01$, *** $p<0.001$, **** $p<0.0001$.

**Table S12.** Statistical differences between SR (%) values obtained after 24 h of incubation of BC samples in water.

| | M1 | M2 | M3 | M4 | M5 | M6 | M7 | C |
|---|---|---|---|---|---|---|---|---|
| M1 | × | * | ns | **** | ns | ns | ** | **** |
| M2 | * | × | *** | ** | ns | ns | ns | **** |
| M3 | ns | *** | × | **** | * | * | **** | **** |
| M4 | **** | ** | **** | × | **** | **** | ** | ns |
| M5 | ns | ns | * | **** | × | ns | ns | **** |
| M6 | ns | ns | * | **** | ns | × | ns | **** |
| M7 | ** | ns | **** | ** | ns | ns | × | *** |
| C | **** | **** | **** | ns | **** | **** | *** | × |

C – control BC sample; M1-M7 – modified BC samples; * $p<0.05$, ** $p<0.01$, *** $p<0.001$, **** $p<0.0001$.

**Table S13.** Statistical differences between WHC (%) values obtained after 60 min of the incubation of BC samples at 37°C.

| | M1 | M2 | M3 | M4 | M5 | M6 | M7 | C |
|---|---|---|---|---|---|---|---|---|
| M1 | × | ns | ns | ns | ns | ns | ns | **** |
| M2 | ns | × | ns | ns | ns | ns | ns | **** |
| M3 | ns | ns | × | ns | ns | ns | ns | **** |
| M4 | ns | ns | ns | × | ns | ns | ns | **** |
| M5 | ns | ns | ns | ns | × | ns | ns | **** |
| M6 | ns | ns | ns | ns | ns | × | ns | **** |
| M7 | ns | ns | ns | ns | ns | ns | × | **** |
| C | **** | **** | **** | **** | **** | **** | **** | × |

C – control BC sample; M1-M7 – modified BC samples; * $p<0.05$, ** $p<0.01$, *** $p<0.001$, **** $p<0.0001$.

**Table S14.** Statistical differences between weight (g) values of BC samples after 30 min of centrifuging (200 *g*).

| | M1 | M2 | M3 | M4 | M5 | M6 | M7 | C |
|---|---|---|---|---|---|---|---|---|
| M1 | × | ns | ns | ns | ns | ns | ns | **** |
| M2 | ns | × | ns | ns | ns | ns | ns | **** |
| M3 | ns | ns | × | ns | ns | ns | ns | **** |
| M4 | ns | ns | ns | × | ns | ns | ns | **** |
| M5 | ns | ns | ns | ns | × | ns | ns | **** |
| M6 | ns | ns | ns | ns | ns | × | ns | **** |
| M7 | ns | ns | ns | ns | ns | ns | × | **** |
| C | **** | **** | **** | **** | **** | **** | **** | × |

C – control BC sample; M1-M7 – modified BC samples; * $p<0.05$, ** $p<0.01$, *** $p<0.001$, **** $p<0.0001$.

**Table S15.** Statistical differences between the values of the density of BC samples.

| | M1 | M2 | M3 | M4 | M5 | M6 | M7 | C |
|---|---|---|---|---|---|---|---|---|
| M1 | × | ns | ns | ** | ns | ns | ns | **** |
| M2 | ns | × | ns | * | ns | ns | ns | **** |
| M3 | ns | ns | × | *** | ns | ns | ns | **** |
| M4 | ** | * | *** | × | ** | * | * | **** |
| M5 | ns | ns | ns | ** | × | ns | ns | **** |
| M6 | ns | ns | ns | * | ns | × | ns | **** |

| | | | | | | | | |
|---|---|---|---|---|---|---|---|---|
| M7 | ns | ns | ns | * | ns | ns | × | **** |
| C | **** | **** | **** | **** | **** | **** | **** | × |

C – control BC sample; M1-M7 – modified BC samples; * p<0.05, ** p<0.01, *** p<0.001, **** p<0.0001.

**Table S16.** Adjusted p-values for the results presented in Table S11 – S15.

| | * | ** | *** | **** |
|---|---|---|---|---|
| SR% after 60 min | 0.0711 – 0.9997 | 0.0149 | 0.0032 – 0.0061 | < 0.0001 |
| SR% after 24 h | 0.0107 – 0.0254 | 0.0018 – 0.0074 | 0.0002 – 0.0003 | < 0.0001 |
| WHC % after 60 min | - | - | - | < 0.0001 |
| Weight (g) after 30 min of centrifuging | - | - | - | < 0.0001 |
| Density | 0.0176 – 0.0215 | 0.0028 – 0.0090 | 0.0006 | < 0.0001 |

### 5. Assessment of cytotoxicity of modified BC

#### *5.1. Extract assay*

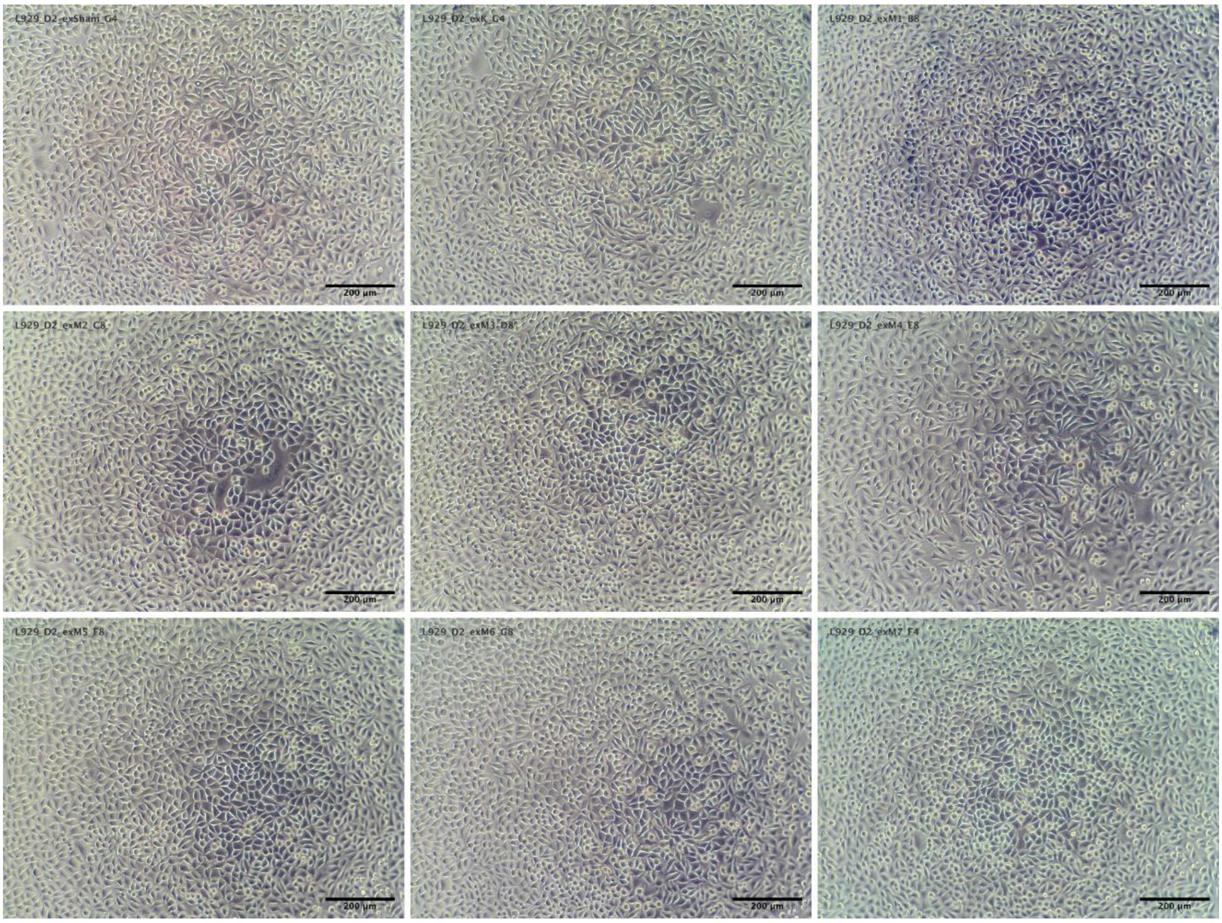


**Figure S11.** Representative micrographs of L929 cells after 24 h of culture with BC extracts. Sequentially from top left to bottom right: sham, control BC sample, M1-M7 – modified BC samples. Scale bar represents 200 µm.

#### 5.2. Direct contact assay

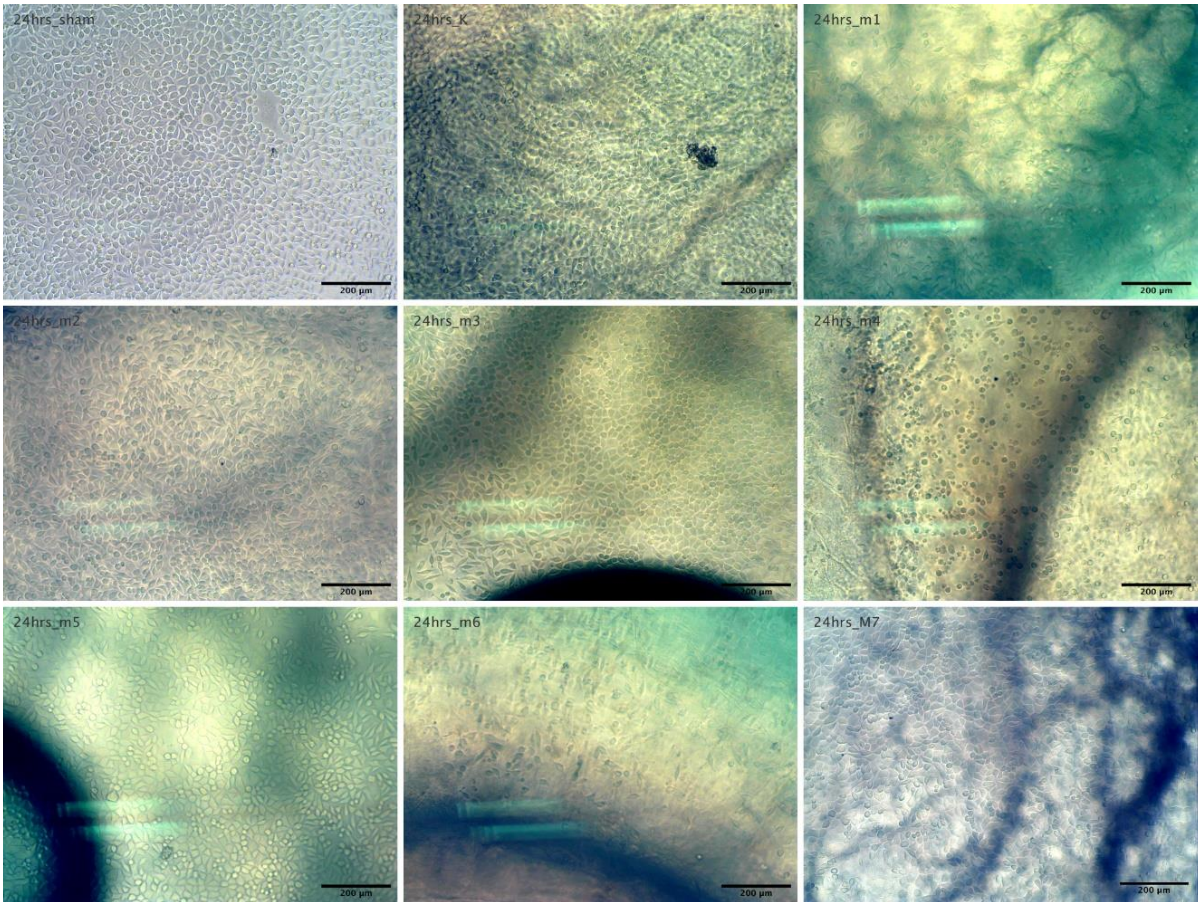

**Figure S12.** Representative micrographs of L929 cells after 24 h of culture beneath sham (CellCrown insert alone, top left panel) and cellulose discs; sequentially from top left to bottom right: control BC sample, M1-M7 – modified BC samples. Scale bar represents 200 µm. Overall, the CellCrown insert made visualization at the edges of the well difficult, thus only the center was imaged. To avoid any disruption to the monolayer, discs were not removed, thus image quality is reduced due to the effect of the sample on the transmitted light.

**Table S17.** Statistical differences between normalized cell viability (% of Sham) in extract assay.

| | M1 | M2 | M3 | M4 | M5 | M6 | M7 | C |
|---|---|---|---|---|---|---|---|---|
| M1 | × | ns | ns | * | ns | ns | ** | **** |
| M2 | ns | × | ** | **** | ns | * | ns | **** |
| M3 | ns | ** | × | ns | ns | ns | **** | **** |
| M4 | * | **** | ns | × | * | ns | **** | **** |
| M5 | ns | ns | ns | * | × | ns | ** | **** |
| M6 | ns | * | ns | ns | ns | × | *** | **** |
| M7 | ** | ns | **** | **** | ** | *** | × | **** |
| C | **** | **** | **** | **** | **** | **** | **** | × |

C – control BC sample; M1-M7 – modified BC samples; * p<0.05, ** p<0.01, *** p<0.001, **** p<0.0001.

**Table S18.** Statistical differences between normalized cell viability (% of Sham) in direct contact assay.

| | M1 | M2 | M3 | M4 | M5 | M6 | M7 | C |
|---|---|---|---|---|---|---|---|---|
| M1 | × | ns | ns | * | ns | ns | ns | ns |
| M2 | ns | × | ns | * | ns | ns | ns | ns |
| M3 | ns | ns | × | ns | ns | ns | ** | ns |
| M4 | * | * | ns | × | ns | ** | **** | *** |
| M5 | ns | ns | ns | ns | × | ns | ** | ns |
| M6 | ns | ns | ns | ** | ns | × | ns | ns |
| M7 | ns | ns | ** | **** | ** | ns | × | ns |
| C | ns | ns | ns | *** | ns | ns | ns | × |

C – control BC sample; M1-M7 – modified BC samples; * p<0.05, ** p<0.01, *** p<0.001, **** p<0.0001.

**Table S19.** Adjusted p-values for the results presented in Table S17 – S18.

| | * | ** | *** | **** |
|---|---|---|---|---|
| Contact assay | 0.0140 – 0.0198 | 0.0045 – 0.0070 | 0.0005 | < 0.0001 |
| Extract assay | 0.0173 – 0.0231 | 0.0017 – 0.0074 | 0.0002 | < 0.0001 |

### 6. Comparison of swelling ability of modified BC with commercial superabsorbent dressings

**Table S20.** Comparison of SR (%) values after 24 h for commercial dressings and M3 sample.

| | SR(%) after 24 h | |
|---|---|---|
| Polyacrylate fiber superabsorbent commercial dressing | 1432.23 ± 37.25 | 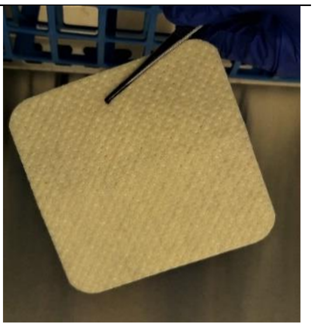 |
| Hydrofiber superabsorbent commercial dressing | 2590.53 ± 235.33 | 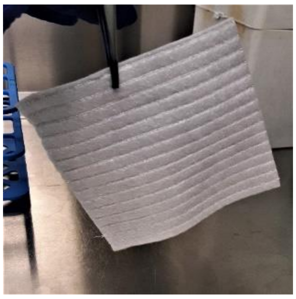 |
| Modified BC (M3) | 3334.21 ± 353.54 | 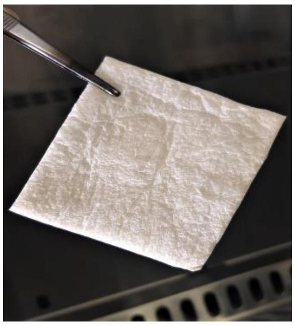 |

**TableS21.** Statistical differences between SR (%) of commercial dressings and M3 sample.

| | Fiber dressing | Hydrofiber dressing | M3 |
|---|---|---|---|
| Fiber dressing | × | ** | *** |
| Hydrofiber dressing | ** | × | * |
| M3 | *** | * | × |

* p<0.05, ** p<0.01, *** p<0.001.

**Table S22.** Adjusted p-values for SR (%) values for the results presented in Table S21.

| | * | ** | *** | **** |
|---|---|---|---|---|
| SR(%) | 0.0319 | 0.0041 | 0.0001 | - |
